# Working memory impairments in Neurofibromatosis Type 1 are explained by disrupted functional connectivity and network controllability

**DOI:** 10.1101/2025.04.10.648210

**Authors:** Marta Czime Litwińczuk, Nelson J Trujillo-Barreto, Jeyoung Jung, Yuping Yang, Shruti Garg, Caroline Lea-Carnall

## Abstract

Neurofibromatosis type 1 (NF1) is a rare, monogenic condition associated with impairments in working memory processing. However, it is unknown to what extent these impairments are explained by disrupted connectivity and state transitions of the functional brain networks. This study characterised task-state functional connectivity (FC) and network controllability during working memory in NF1 and investigated whether altered controllability distinguished NF1 from neurotypical controls and related to individual differences in working memory performance. We analysed functional MRI data from 53 NF1 adolescents and 36 age-matched neurotypical controls performing a verbal N-back task. Compared to controls, NF1 participants showed reduced FC within frontoparietal network (particularly between the left prefrontal cortex and left intraparietal sulcus), reduced FC between default and limbic networks, and increased FC between control and somatomotor regions. Network controllability analysis revealed increased average controllability but reduced modal controllability and activation energy in prefrontal, parietal, temporal and retrosplenial regions in NF1, indicating that in NF1 these regions have enhanced capacity to drive the network toward easy-to-reach states and are readily activated within the current network organisation. The opposite pattern of reduced average controllability but increased modal controllability and activation energy were observed in somatosensory, auditory, extrastriate and temporal-parietal regions, suggesting enhanced capacity of these regions to drive the network towards difficult-to-reach states. These changes accurately predicted NF1 status, with an area under the ROC curve exceeding 80% across all controllability models. Furthermore, average controllability accurately predicted working memory performance in the NF1 group, generalising to out-of-sample data. In particular, left and right lateral ventral prefrontal cortex, left superior parietal lobule, left intraparietal sulcus consistently emerged as predictors of working memory performance across all measures. By demonstrating that altered regional network controllability in NF1 predicts working memory performance, this study provides a novel mechanistic account of the functional network alterations underlying cognitive difficulties in NF1 and establishes a framework for evaluating future therapeutic interventions through their effects on functional brain network control.

## Introduction

Neurofibromatosis Type 1 (NF1) is a rare, single-gene neurodevelopmental condition that affects 1 in 2,700 individuals (Evans et al., 2010). It is caused by mutations in the NF1 gene, which encodes the neurofibromin protein. Animal models suggest that this mutation leads to dysregulated gamma-aminobutyric acid (GABA) signalling (Costa et al., 2002). Human research supports this mechanism, demonstrating altered GABAergic neurotransmission and altered balance between excitatory and inhibitory signalling (excitation/inhibition balance) (Violante et al., 2016; Wild et al., 2026). Affected individuals may present with pigmentary lesions (café-au-lait spots), dermal neurofibromas, skeletal abnormalities, brain and peripheral nerve tumours, learning disabilities, and cognitive and social difficulties (Gutmann et al., 2017; Lehtonen et al., 2012). In particular, working memory difficulties have been frequently reported in NF1 (Lehtonen et al., 2012; Plasschaert et al., 2016; Pobric et al., 2021; Sawyer et al., 2022; Shilyansky et al., 2010). Working memory refers to the ability to temporarily remember and manipulate information for a short period of time (Baddeley, 2003). It is a critical ability for learning and decision making (Baddeley, 2012) and it has been shown to impact academic achievement (Swanson & Alloway, 2012). Therefore, understanding the mechanisms of working memory difficulties in NF1 is an important area of research.

Previous work used functional magnetic resonance imaging (fMRI) to study the brain activation patterns related to working memory in NF1 participants. Shilyansky et al. (2010) found that during performance of visuospatial working memory, NF1 participants show less functional magnetic resonance imaging (fMRI) activation of dorsolateral prefrontal cortex (dlPFC), striatum, frontal eye fields, and parietal cortex compared to neurotypical controls.

These regions are widely known to be active during working memory tasks in neurotypical groups (Owen et al., 2005), which suggests that participants with NF1 may have reduced recruitment of neural resources in brain regions responsible for working memory processes. Ibrahim et al. (2017) focused on studying how working memory affects functional activation and functional connectivity (FC), which reflects the temporal synchrony of activity between pairs of regions, and is thought to reflect neural communication during cognitive processing (Friston, 2011). Ibrahim et al. (2017) found that NF1 participants had greater activation of the posterior cingulate cortex (PCC) and temporal regions, compared to controls. To examine task-related changes in FC, Ibrahim et al. (2017) conducted a psychophysiological interactions analysis. This revealed showed that during working memory NF1 participants had greater connectivity between visual cortices and both PCC and parietal regions, as well as between PCC and the cerebellum, as compared to a control group. This suggests that the NF1 group recruited different neural resources to achieve task performance. Importantly, PCC is usually deactivated during task performance and it is active during rest along with other regions belonging to the default mode network (DMN) (Raichle et al., 2001). The authors therefore considered whether the NF1 group’s brain activity patterns may be explained by the “DMN interference” hypothesis (Violante et al., 2012). According to this hypothesis, overactivation of the DMN during task performance may interfere with network activity directly related to the task. This provides a potential explanation for the altered patterns of brain activity observed in NF1 during working memory, both at the level of large-scale networks and individual brain regions.

One region of particular importance to working memory is the dlPFC (Barbey et al., 2013; Owen et al., 2005). In NF1, both fMRI and electroencephalography studies have shown hypoactivation of the left dlPFC during working memory tasks (Ibrahim et al., 2017; Pobric et al., 2021). This motivated investigations into whether non-invasive brain stimulation (NIBS) can be used as a therapeutic tool to directly modulate dlPFC activity and improve working memory performance in NF1 (Garg et al., 2022). Researchers administered anodal transcranial direct current stimulation (tDCS) to the left dlPFC of adolescents with NF1 to test the hypothesis that increasing cortical excitability and reducing local GABA concentrations would enhance cognitive performance during a verbal working memory task. It was found that anodal tDCS successfully maintained dlPFC activity levels from baseline throughout the task, whereas during sham stimulation dlPFC activity decreased as the task progressed. However, these physiological shifts did not produce the predicted behavioural gains; working memory performance worsened during stimulation, and no significant improvements were observed at the end of the intervention. (Garg et al., 2022). Further analysis of this dataset showed that anodal tDCS altered dlPFC’s effective connectivity, with decreased connectivity within frontal cortex and increased connectivity within striatum after stimulation (Litwińczuk et al., 2025). Together, these results show that tDCS can modify both regional activity and the connectivity of the stimulated region.

However, the lack of cognitive improvement after tDCS suggests that local stimulation of task-activated regions may not be sufficient to overcome the broader network-level disruptions in NF1. Currently, brain regions targeted by brain stimulation approaches are largely determined based on fMRI task activation maps. However, these maps often fail to capture the underlying network dynamics that allow a region to influence broader cognitive states. This may limit our ability to identify the mechanisms of cognitive impairment in NF1, as regional activity levels do not necessarily reflect a node’s capacity to drive coordinated activity across the connectome. Therefore, examining how NF1 alters the ability of specific brain regions to influence whole-brain dynamics may provide greater insight into the neural mechanisms underlying working memory deficits, and aid identification of brain stimulation targets.

Network control theory provides a suitable framework for this type of characterisation by estimating how effectively each brain region can drive the network to transition between states (Bassett et al., 2011; Gu et al., 2015). Within this framework, brain states represent specific patterns of underlying neural activity (Bassett et al., 2011). Network controllability theory provides metrics, such as average and modal controllability, that can be used to characterise brain regions in terms of their ability to drive the system towards easy-to-reach states (low energy demanding) and difficult-to-reach states (high energy demanding), respectively. Importantly, it has been shown that the type of controllability associated with each area is related to their connectivity degree. For example, research in neurotypical cohorts has shown that densely connected regions, particularly parts of the default mode network, are associated with high average controllability (Gu et al., 2015). Meanwhile, weakly connected (peripheral) regions, particularly in the executive control network that is typically active during working memory, were associated with high modal controllability (Gu et al., 2015; Saberi et al., 2024). Beynel et al. (2020) provide a proof of concept for the functional relevance of the network controllability framework. In their study, brain stimulation targets for young, neurotypical adults were identified based on task-related fMRI activation, and subsequent analyses showed that the modal controllability of these regions predicted the degree of stimulation-induced improvement in working memory performance. Specifically, stimulation of regions with high task activation and high modal controllability resulted in the greatest behavioural gains. This demonstrates that controllability metrics capture the propensity of a region to facilitate task-relevant state transitions. The present study will build on this by investigating whether altered controllability profiles themselves can serve as markers for working memory impairment in NF1, shifting the focus from regional activation levels to the underlying control capacity of the network.

Network controllability is typically estimated from structural connectivity (SC) derived from diffusion tensor imaging, providing a map of the physical white matter pathways that constrain the system’s transition capacity. However, there is an increasing shift in research toward applying these metrics to functional brain networks (Wang et al., 2025; J. Yang et al., 2025; Y. Yang et al., 2025), which describe the patterns of coordinated activity between brain regions. While SC-based controllability reflects the brain’s anatomical architecture, FC-based controllability provides a window into the dynamic, task-dependent ’state-space’ and the network’s operational ability to navigate it in real-time. Recent work demonstrates that these functional control profiles can reveal network alterations that remain hidden in standard connectivity analyses. For example, J. Yang et al. (2025) identified shared patterns of frontoparietal dysconnectivity across psychiatric conditions like schizophrenia, bipolar disorder, and major depressive disorder. Crucially, by applying controllability analysis to these functional networks, they were able to move beyond simply identifying disconnections to reveal disorder-specific impairments in the network’s ability to drive transitions across states during cognitive processing. This provides a rationale for applying controllability to functional brain networks when investigating working memory in NF1.

The present work investigates *i*) how FC and controllability are altered in NF1 during working memory performance, *ii*) whether disrupted control profiles can serve as markers of NF1 and *iii*) how they relate to working memory performance. To achieve these aims, we examined functional brain organisation during the 2-back (working memory) condition of a verbal N-back in 53 adolescents diagnosed with NF1 and 36 age-matched neurotypical controls. First, we compared FC and network controllability measures between the groups to identify alterations associated with NF1. Next, we examined whether regions showing altered controllability could distinguish individuals with and without an NF1 diagnosis, and whether these alterations were related to individual differences in working memory performance within the NF1 group.

We hypothesise that *i*) compared to neurotypical controls, the NF1 group will show increased connectivity between the default mode and frontoparietal networks; *ii*) the network architecture in NF1 will show a reduced capacity to drive the system towards difficult-to-reach states within executive regions (reduced modal controllability), and *iii*) following the ‘default mode interference’ hypothesis hubs within the default mode network will exhibit an increased bias towards easy-to-reach states (increased average controllability); and *iv*) these altered control profiles will be predictive of poorer working memory performance in NF1 participants.

## Results

### NF1 group shows working memory impairment that increases with cognitive load

We first assessed working memory performance during the in-scanner verbal N-back task to establish whether NF1 participants showed behavioural impairments (Figure 1), providing a behavioural context for subsequent FC and network controllability analyses. The group differences during out-of-scanner cognitive assessments (visuospatial N-back and Corsi Blocks) are reported in Supplementary Material 1.

**Figure 1.**
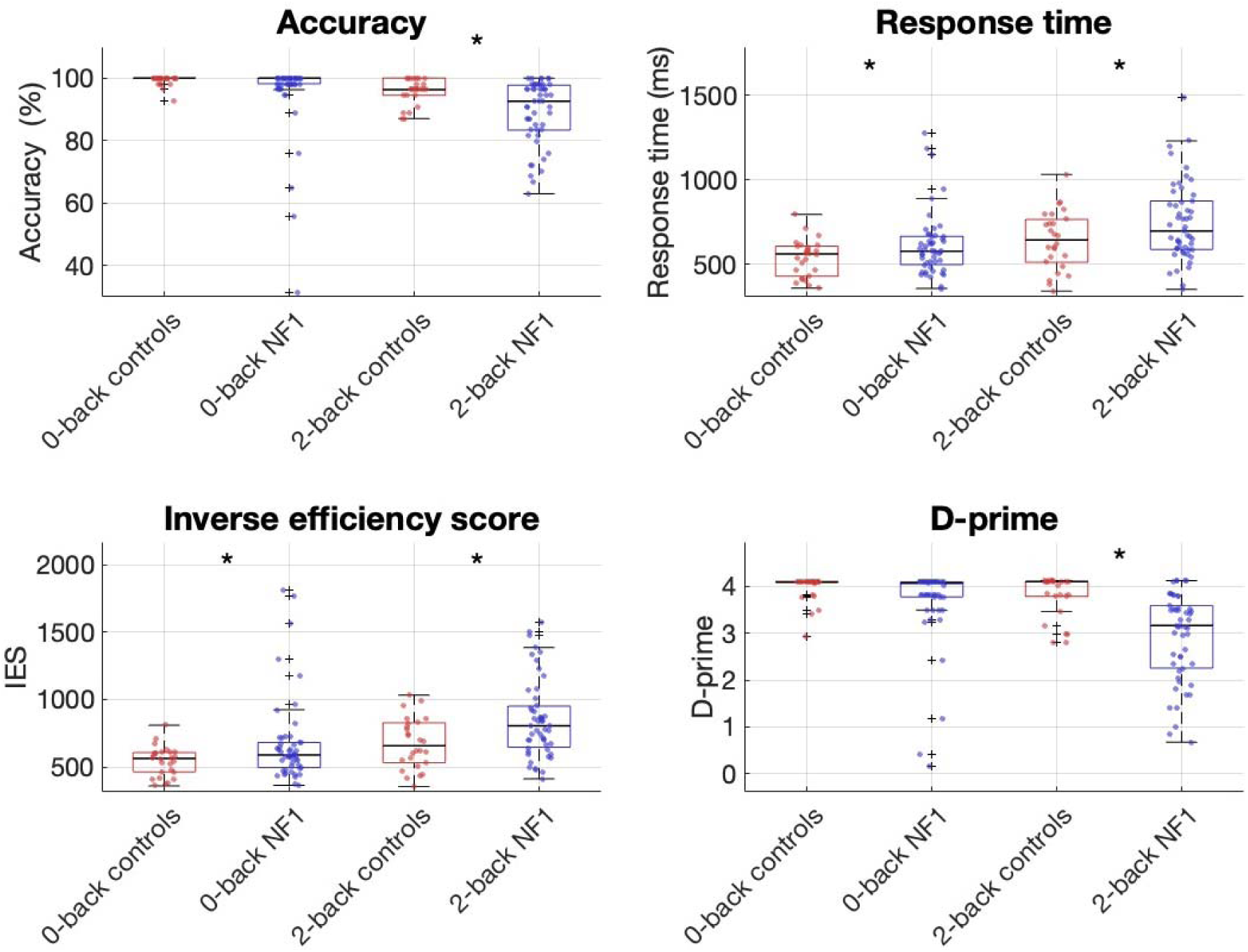
Boxplots illustrate NF1 participants’ and controls’ in-scanner N-back task performance. Boxes represent the interquartile range (IQR) containing the middle 50% of the data, the black horizontal lines inside indicate the median, the whiskers extend to the most extreme data not considered outlier. Individual dots represent individual participants’ scores. Stars indicate significant differences between NF1 participants and controls, not corrected for multiple comparisons.

During the 0-back condition, the NF1 group showed slower response times than controls (*t*(73) = 2.18, *p* = 0.032, β = 85.281), and higher IES (*t*(73) = 2.22, *p* = 0.030, β = 142.485). This impairment was more pronounced during the 2-back condition, in which the NF1 group showed significantly lower accuracy (*t*(73) = -3.06, *p* = 0.003, β = -6.438), slower response times (*t*(73) = 2.24, *p* = 0.028, β = 111.392), higher IES (*t*(73) = 3.02, *p* = 0.003, β = 183.628), and lower d-prime (*t*(73) = -4.46, *p* < 0.001, β = -0.883) than the control group. Importantly, the 2-back>0-back contrast revealed that the NF1 group showed significantly greater decline in d-prime (*t*(73) = -2.87, *p* = 0.005, β = -0.523).

Together, these findings indicate that the NF1 group showed poorer working memory performance, with greater decline in performance as cognitive load increased.

### Functional connectivity is reduced in higher-order associative networks but increased in somatomotor and medial prefrontal cortex in NF1

Next, we tested whether the behavioural differences in working memory performance in NF1 were accompanied by altered FC during the working memory task. Based on our hypothesis that NF1 would show increased connectivity between the default mode and frontoparietal networks, we examined whole-brain FC during the 2-back condition (Figure 2; Supplementary Tables 1 and 2). The 2-back FC results revealed three overarching patterns of altered connectivity: i) reduced connectivity affecting higher-order associative networks, including control, default and limbic networks; ii) increased connectivity involving somatomotor, attention and medial prefrontal regions; and iii) a complex pattern of altered subcortical connectivity.

**Figure 2.**
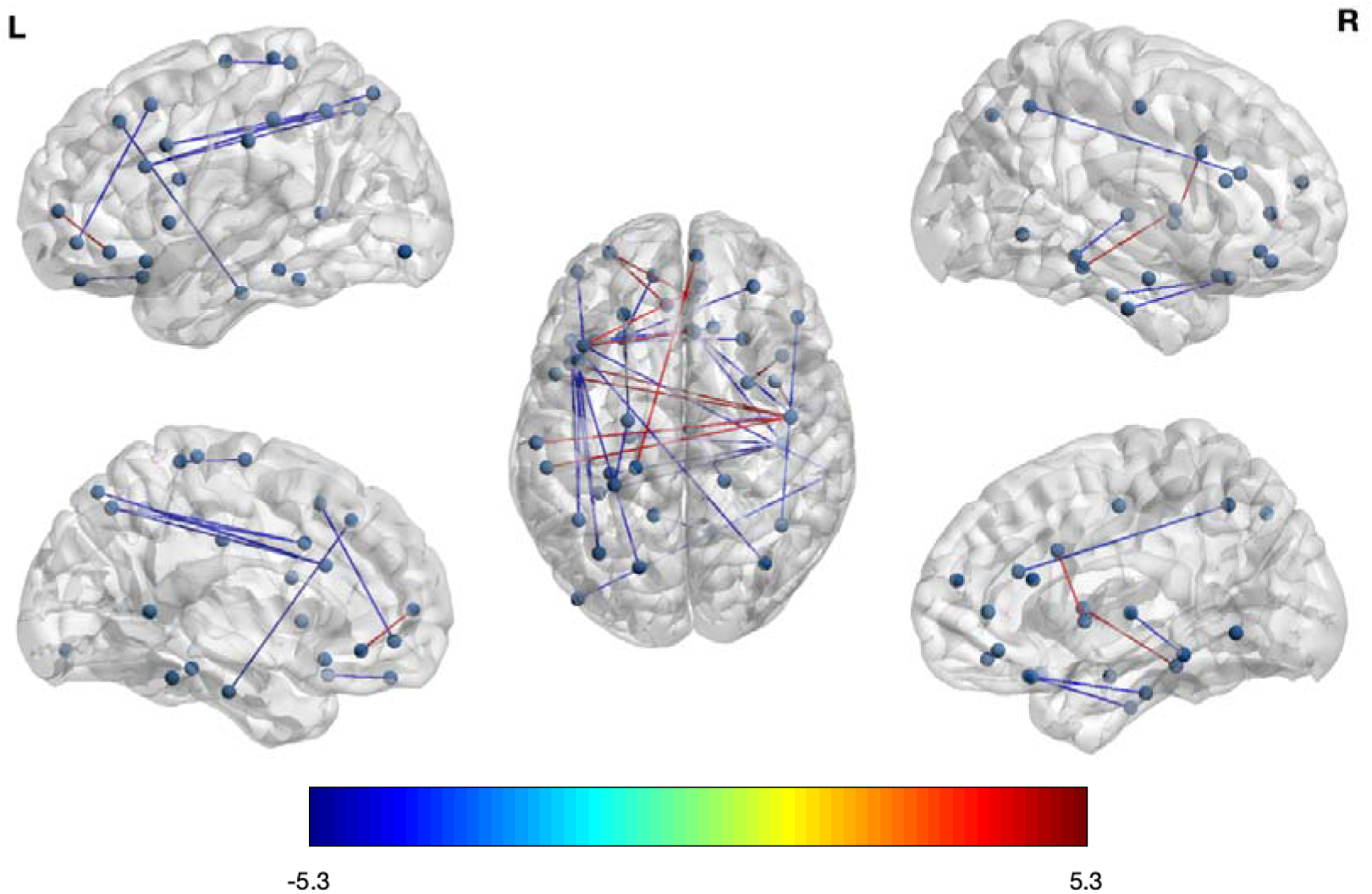
Whole brain FC differences between controls and NF1 participants during the 2-back condition. Red lines indicate stronger connections in NF1 participants, whereas blue lines indicate stronger connections in controls. Scale indicates t-values. All t-values have passed the probability threshold of 0.05 (FDR-corrected).

**Table 1.** Demographic data of study participants (53 participants with NF1, 36 neurotypical controls).

|  | Mean Age (STD) | Age range | Sex (M/F) |
| --- | --- | --- | --- |
| NF1 participants | 14.36 (1.95) | 11.0-18.3 | 26/27 |
| Controls | 14.29 (2.61) | 6.1-17.9 | 19/17 |

**Table 2.** Classification performance summary for all models using region-wise controllability metrics showing significant group differences in classification analysis. All classification performance metrics are expressed in percentages (%). Values represent mean ± standard deviations across 5,000 evaluations (1,000 repetitions × 5-fold cross-validation). All models included age and sex as covariates.

| **Metric** | **Average**<br>**controllability** | **Modal**<br>**controllability** | **Activation**<br>**energy** | **Combined** |
| --- | --- | --- | --- | --- |
| Accuracy | 77.6 ± 10.4 | 75.1 ± 9.0 | 74.6 ± 9.8 | 77.8 ± 8.7 |
| Balanced<br>Accuracy | 76.9 ± 10.9 | 74.1 ± 9.5 | 73.3 ± 10.3 | 76.9 ± 9.2 |
| Sensitivity | 80.8 ± 12.8 | 79.2 ± 12.4 | 80.4 ± 12.4 | 81.8 ± 11.6 |
| Specificity | 73.0 ± 17.5 | 69.0 ± 17.6 | 66.2 ± 17.7 | 72.0 ± 16.5 |
| Precision | 82.4 ± 10.3 | 80.0 ± 9.6 | 78.6 ± 9.6 | 82.0 ± 9.2 |
| AUC | 85.5 ± 9.1 | 82.7 ± 9.2 | 82.9 ± 9.4 | 85.4 ± 8.1 |

### NF1 shows reduced connectivity within and between control, default and limbic networks, particularly in left prefrontal, parahippocampal and temporo-occipital cortices

Reductions in FC were most prominent within the control network, with lateral prefrontal cortex showing reduced connectivity with the intraparietal sulcus and inferior parietal lobule bilaterally, both within and across hemispheres. The control network also had reduced connectivity with default, dorsal attention, limbic, and central visual networks. Of the control B network, left lateral ventral prefrontal cortex and right temporal cortex showed reduced connectivity with left lateral prefrontal cortex (default B); right lateral ventral prefrontal cortex showed reduced connectivity with left orbital frontal cortex (limbic B); right temporal cortex showed reduced connectivity with left extrastriate cortex (central visual). Additionally, left lateral prefrontal cortex (control A) showed reduced connectivity with left superior parietal lobule (dorsal attention A).

Within the default network, reduced connectivity was found between left dorsal prefrontal cortex and left parahippocampal cortex, and between left parahippocampal cortex and right temporal cortex. The default network also had reduced connectivity with the limbic network; between left ventral prefrontal cortex (default B) and right orbital frontal cortex (limbic B), between left parahippocampal cortex (default C) and right temporal pole (limbic A). Finally, default network connectivity reductions were found between left retrosplenial cortex (default C) and right extra-striate inferior cortex (peripheral visual).

Within the limbic network, reduced connectivity was observed between bilateral orbital frontal cortex regions and between right orbital frontal cortex and right temporal pole. Additionally, left temporal occipital cortex (dorsal attention A) had reduced connectivity with right temporal pole (limbic A).

Overall, the reductions in NF1 during working memory condition largely affected the left hemisphere, involving left lateral, ventral and dorsal prefrontal, parahippocampal, and temporal occipital cortices.

### NF1 shows increased connectivity involving right somatomotor and medial prefrontal cortex, extending to left-hemisphere control, default, attention and somatomotor networks

The NF1 participants showed increased connectivity with right somatomotor cortex (somatomotor A) and right medial prefrontal cortex (default A). Then, in NF1 right somatomotor cortex showed increased connectivity left lateral prefrontal cortex (control A), left postcentral cortex (dorsal attention B) and left insula (ventral attention B). Similarly, right medial prefrontal cortex showed increased connectivity with left lateral ventral prefrontal cortex (control B), left lateral prefrontal cortex (default B) and left somatomotor cortex (somatomotor A). Contralaterally, left medial prefrontal cortex (default A) also showed increased connectivity with left lateral ventral prefrontal cortex (control B), and left lateral prefrontal cortex (default B). Additionally, right insula (ventral attention A) showed increased connectivity with right temporal cortex (control B).

Overall, connectivity increases in NF1 during working memory condition were right-lateralised, with right somatomotor and right medial prefrontal cortex both showing stronger connectivity to left hemisphere control, default, dorsal and ventral attention, and somatomotor networks.

### NF1 shows increased prefrontal-striatal connectivity but decreased visual-thalamic connectivity

Subcortical connectivity showed a complex pattern in NF1: right lateral prefrontal cortex (control A) showed increased connectivity with the right putamen, while right extra-striate inferior cortex (peripheral visual) showed reduced connectivity with the right mediodorsal thalamus.

### Network-level analyses confirm reductions within and between control, limbic and default networks in NF1 during working memory task, with modest increases involving subcortical-limbic and somatomotor-temporal parietal connections

After examining region-wise group differences, we examined whether these differences reflected broader alterations in connectivity between canonical functional systems (Yeo et al., 2011) by averaging connection strength across all connections within each network and between each pair of networks. This network-level analysis was consistent with the region-wise findings reported above, but these findings did not survive FDR correction for multiple comparisons (Figure 3).

**Figure 3.**
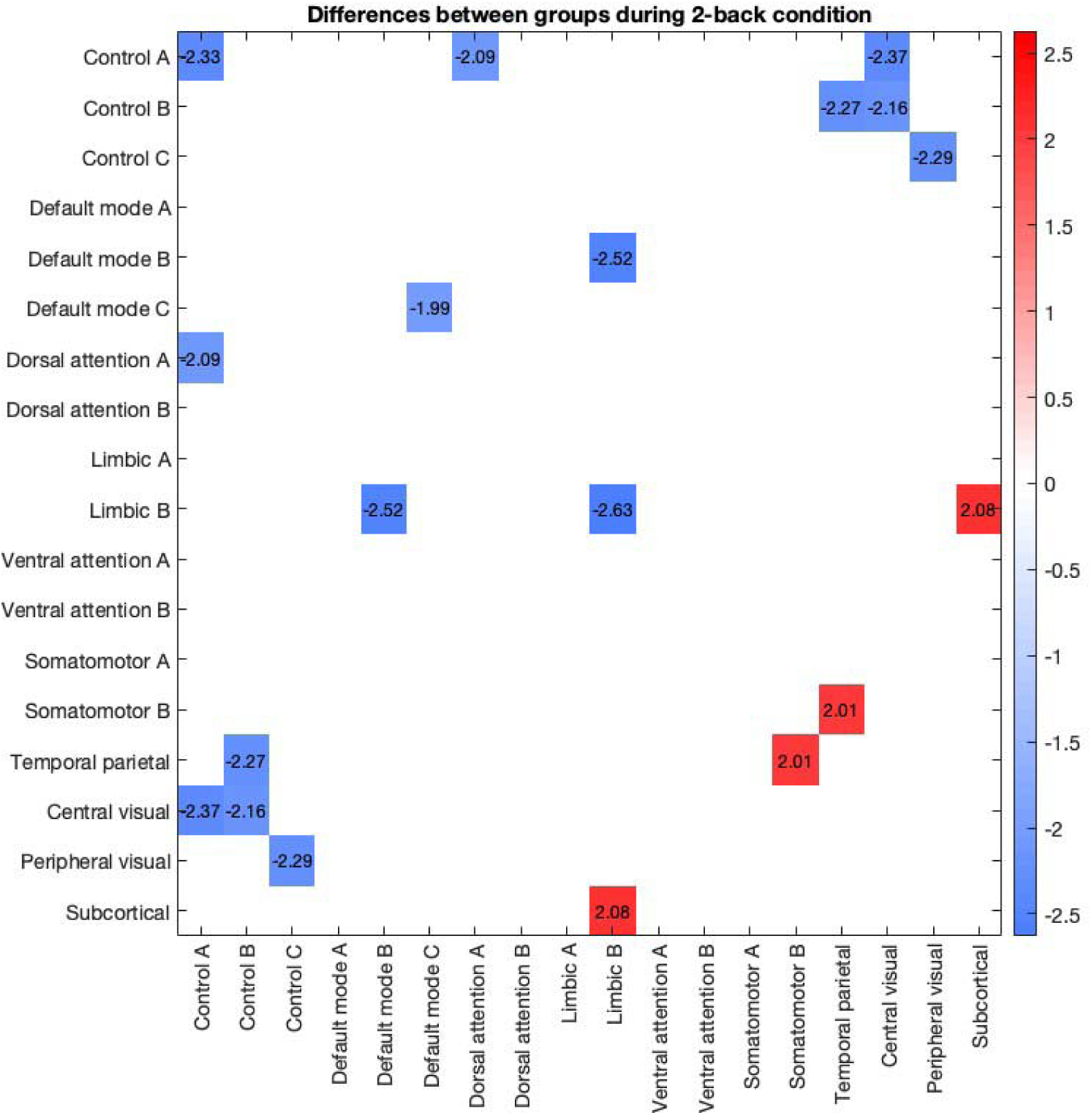
Group differences between NF1 participants and controls in functional brain networks during 2-back condition. Values in the heatmap represent t-statistics. Red values indicate stronger connectivity for NF1 participants and blue values indicate weaker connectivity in NF1 participants. All t-values have passed uncorrected p-value threshold of 0.05.

Specifically, NF1 participants showed reduced within-network connectivity in control A (*t* = −2.33, *p* = 0.022), limbic B (*t* = −2.63, *p* = 0.010), and default C (*t* = −1.99, *p* = 0.050). Reduced between-network connectivity was observed for limbic B and default B (*t* = −2.52, *p* = 0.014), central visual and control A (*t* = −2.37, *p* = 0.020), central visual and control B (t = −2.16, *p* = 0.033), peripheral visual and control C (*t* = −2.29, *p* = 0.024), temporal parietal and control B (*t* = −2.27, *p* = 0.026), and dorsal attention and control A (*t* = −2.09, *p* = 0.039) networks. Conversely, NF1 participants showed increased connectivity between subcortical and limbic B networks (*t* = 2.08, *p* = 0.041), and temporal parietal and somatomotor B networks (*t* = 2.01, *p* = 0.048).

### Neurotypical control and NF1 groups show hierarchical organisation of network controllability across the brain

Having established altered FC in NF1 during working memory task, we next investigated whether these changes were associated with altered capacity of brain regions to control network dynamics. We estimated average controllability, modal controllability and activation energy during 2-back condition, and we examined the spatial organisation of controllability measures across the whole brain in each group. Figures 4 and 5 show the regional distributions of average controllability, modal controllability and activation energy in control and NF1 groups, respectively. For neurotypical controls, average controllability showed a hierarchical organisation, with lowest values observed for sensory-motor cortices, higher values observed for higher-order association cortices, and highest values observed for subcortical structures. Within association cortex, moderate values were observed in lateral parietal, posterior frontal, dorsolateral and ventrolateral prefrontal, insular, and lateral temporal regions, whereas the highest cortical values were observed in medial and anterior temporal regions, posterior cingulate and retrosplenial cortex, medial prefrontal cortex, and temporal pole. Among subcortical regions, highest values were observed for thalamic nuclei, locus coeruleus, raphe nuclei, and caudate. The opposite pattern was observed for modal controllability. Activation energy showed lower values in subcortical structures, moderate values for sensory-motor cortices, and highest values for higher-order association cortices.

**Figure 4.**
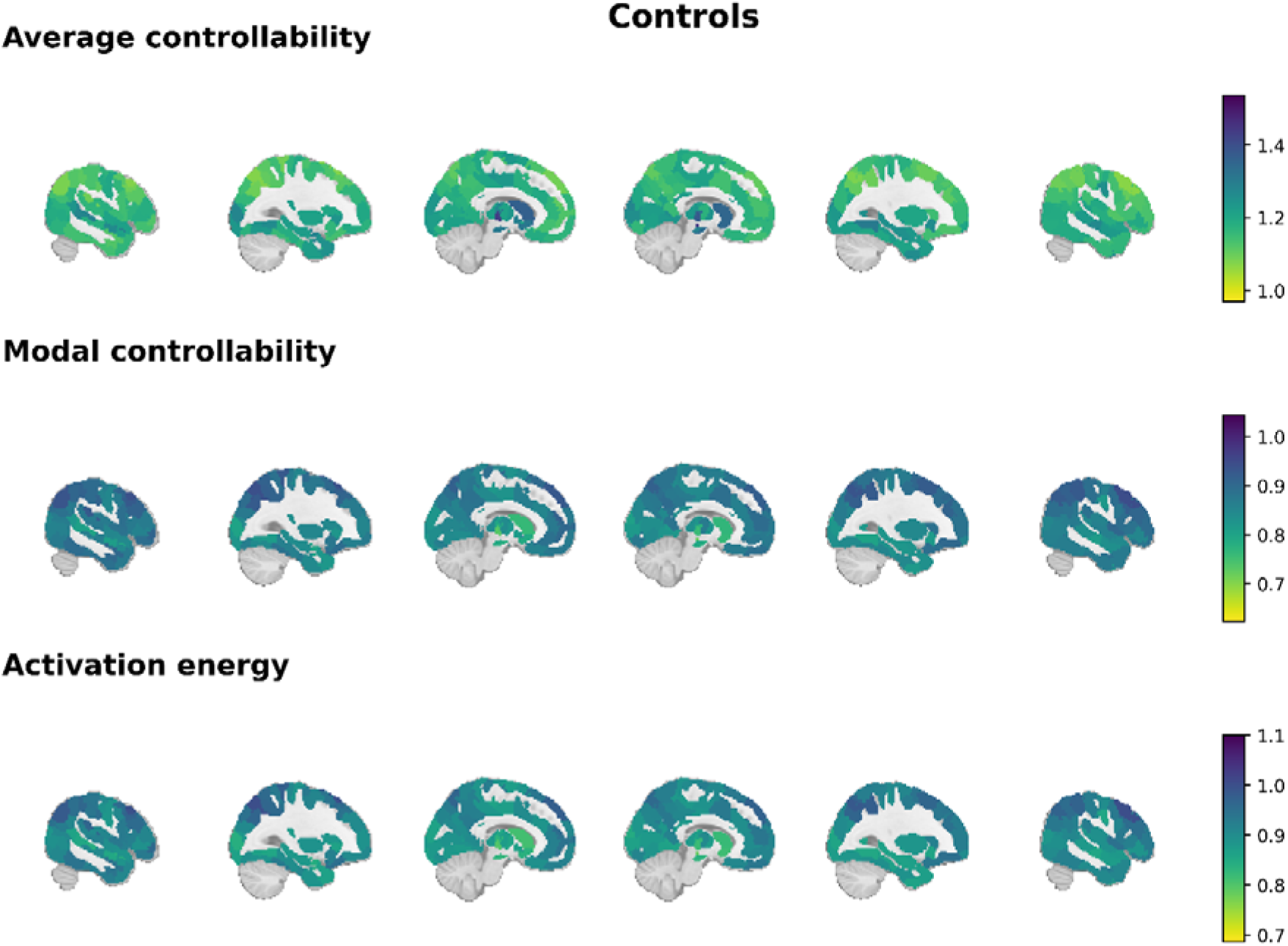
Region-wise network controllability measures in the control group. Maps were generated using the Schaefer 300-region cortical atlas combined with AA3 subcortical regions (354 regions total). Sagittal slices are displayed at coordinates -50,-30,-10, 10, 30, 50, with colour scales representing the magnitude of each network controllability measure. Colour limits are preserved between control and NF1 group for each measure.

**Figure 5.**
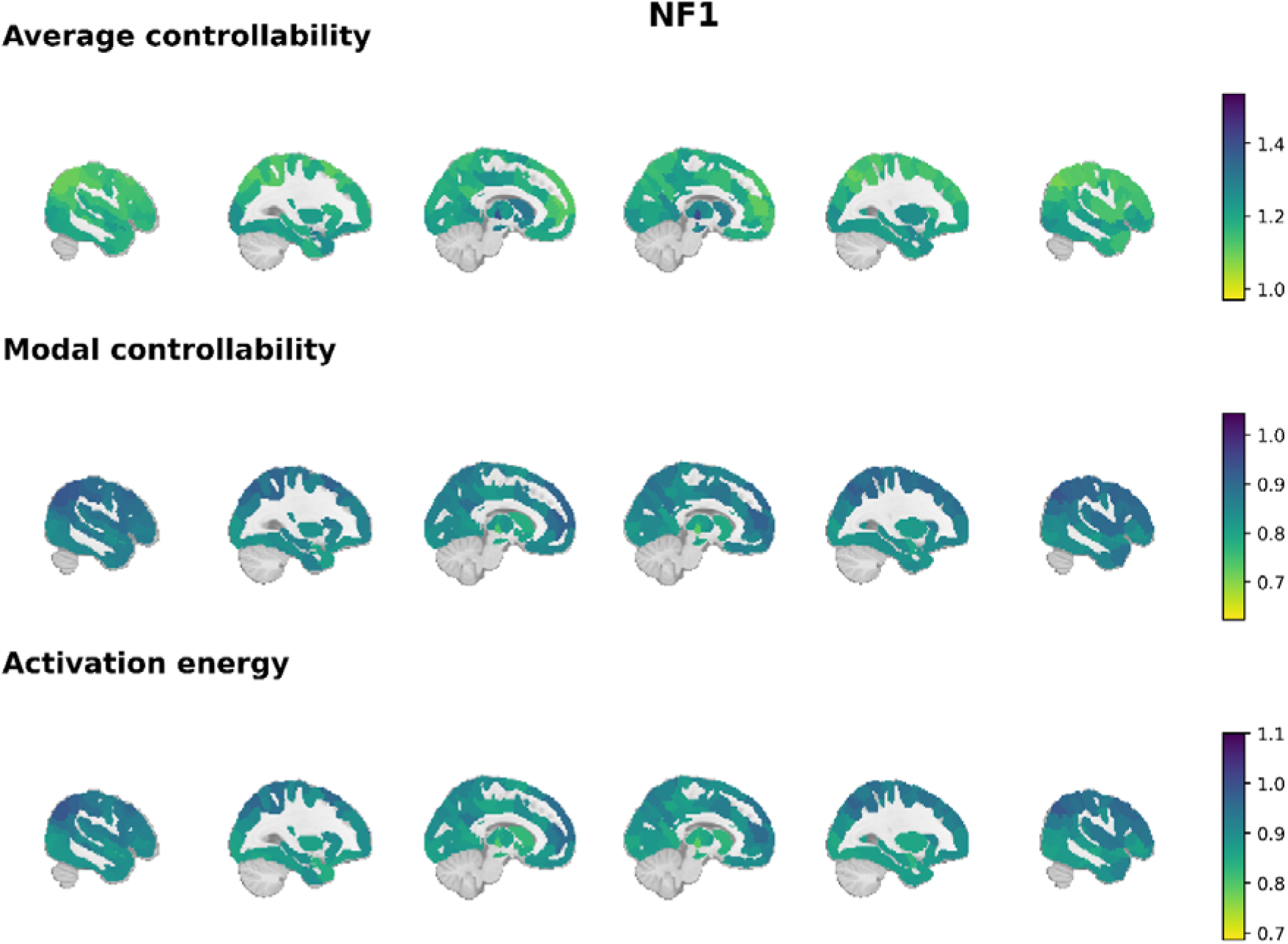
Region-wise network controllability measures in the NF1 group. Maps were generated using the Schaefer 300-region cortical atlas combined with AA3 subcortical regions (354 regions total). Sagittal slices are displayed at coordinates -50,-30,-10, 10, 30, 50, with colour scales representing the magnitude of each network controllability measure. Colour limits are preserved between control and NF1 group for each measure.

The lowest energy values were observed in thalamic and brainstem nuclei, followed by sensory regions, particularly visual cortex. In contrast, the highest activation energy values were observed across higher-order association cortices, including lateral parietal, prefrontal, and temporal association regions. These spatial patterns were broadly preserved in the NF1 group.

### At the network level, NF1 shows increased average controllability but reduced modal controllability and activation energy in dorsal attention, control and default-mode networks, with the opposite pattern in limbic and temporal-parietal networks

After investigating region-wise maps in each group, we investigated whether these spatial patterns observed during working memory task were associated with systematic group differences at the level canonical functional systems (Yeo et al., 2011). The following effects are reported at the uncorrected probability threshold (α = .05). For the results that survive the FDR correction, both the uncorrected p-values and the adjusted q-values are reported.

Specifically, NF1 participants showed reduced modal controllability and activation energy for dorsal attention network A which survived FDR-correction (average controllability: *t* = 2.69, *p* = .009; modal controllability: *t* = -3.09, *p* = .002, *q* = .041; activation energy: *t* = -3.38, *p* = .001, *q* = .020). The same direction of effects was observed at the uncorrected threshold for control network A (average controllability: *t* = 2.03, *p* = .049; modal controllability: *t* = -2.12, *p* = .038; activation energy: *t* = -1.99, *p* = .048), control network B (average controllability: *t* = 2.15, *p* = .030; modal controllability: *t* = -2.24, *p* = .024; activation energy: *t* = -2.21, *p* = .033), and default network C (*t* = 2.36, *p* = .019; modal controllability: *t* = -2.43, *p* = .018; activation energy: *t* = -2.61, *p* = .011).

The opposite pattern of decreased average controllability but increased modal controllability and activation energy in NF1 group, as compared to control group, was seen for the limbic network B (modal controllability: *t* = -1.99, *p* = .048; activation energy: *t* = - 2.04, *p* = .048) and temporal parietal network (average controllability: *t* = -2.24, *p* = .027; modal controllability: *t* = 2.11, *p* = .041).

### At the region level, NF1 shows increased average but reduced modal controllability and activation energy across prefrontal, parietal, temporal and retrosplenial regions, with the opposite pattern in extrastriate, somatomotor, auditory and temporal-parietal regions

Having identified group differences in network controllability during working memory task at the level of canonical functional networks, we next examined which regional differences contributed to these network-level patterns. The following results passed the permutation-based *p*-value threshold of 0.05, without correction for multiple comparisons.

#### Average controllability

The precise region-specific t-statistics and p-values for average controllability are reported in full in Supplementary Table 3 and Figure 6 presents the projection of these results to the brain template. NF1 participants showed higher average controllability than control participants across frontal and parietal regions of control and default mode networks, including the intraparietal sulcus, inferior parietal lobule precuneus lateral ventral and lateral dorsal prefrontal cortex (control network), ventral and medial prefrontal cortex (default mode network), along with the temporal regions (control and default mode) and retrosplenial cortex (default mode network). Beyond control and default mode networks, there was also increased average controllability for NF1 group in superior parietal lobule and frontal eye fields (dorsal attention network), and orbitofrontal cortex (limbic network), lateral prefrontal cortex and parietal operculum (ventral attention network), extrastriate cortex (central visual network network), and ventral posterolateral thalamus (subcortical). In contrast, decreased average controllability in the NF1 group was found for sensory and lower-order networks, including parts of the central visual network (extrastriate cortex), somatosensory and auditory cortices (somatomotor network), and the temporal parietal network.

**Figure 6.**
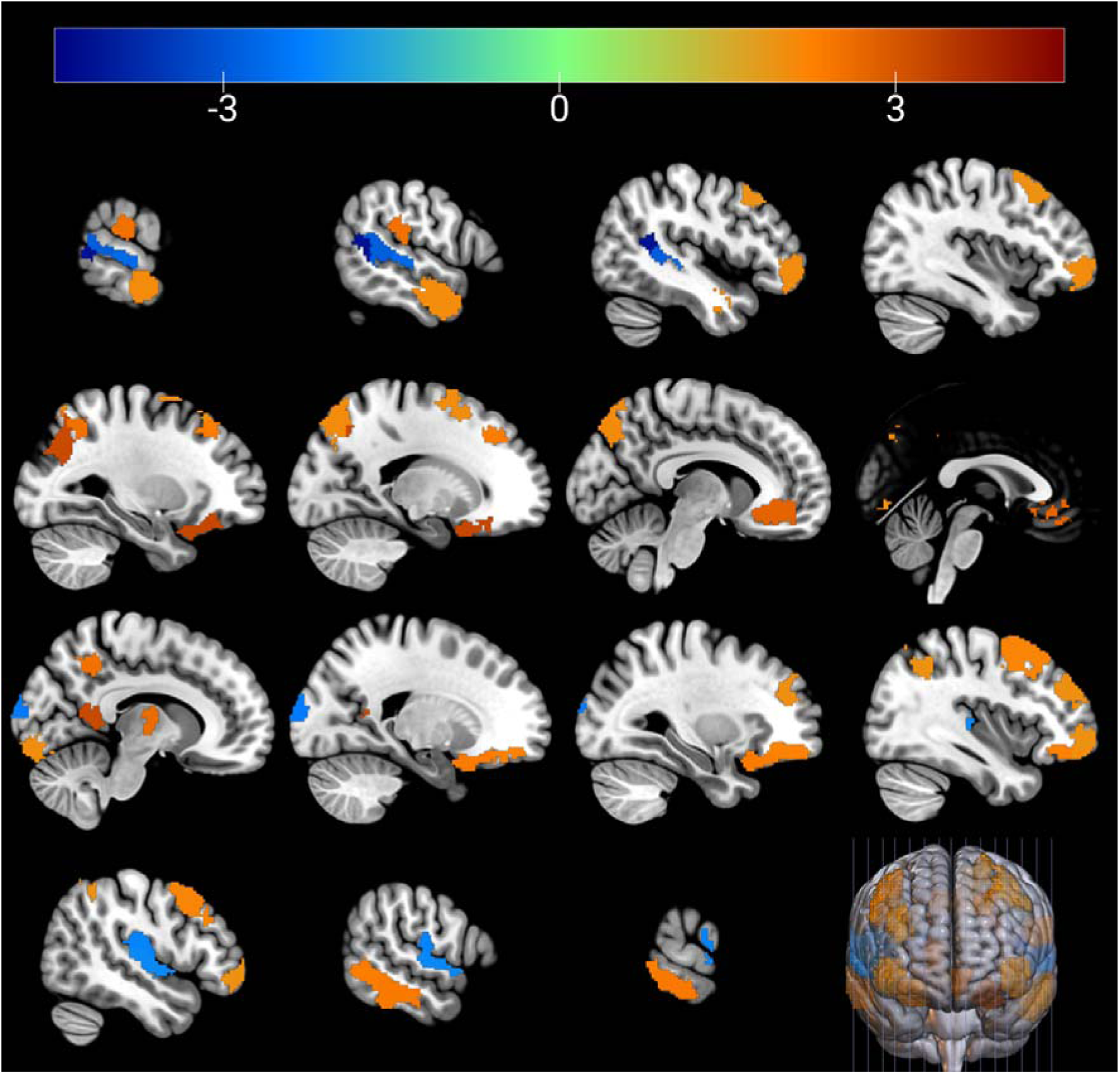
Group differences in average controllability during 2-back task. ROIs are coloured by corresponding t-values (scale: -4.5 to 4.5), where warm colours indicate higher values in NF1, and blue colours indicate lower values in NF1. Sagittal have coordinates: -66, -56, -46, -38; -28, -18, -8, 0; 10, 20, 30, 38; 48, 58, 68.

#### Modal controllability

The region-specific t-statistics and p-values for modal controllability are reported in full in Supplementary Table 4. Figure 7 presents the projection of these results to the brain template. To briefly summarise, the modal controllability findings largely mirror the average controllability findings.

**Figure 7.**
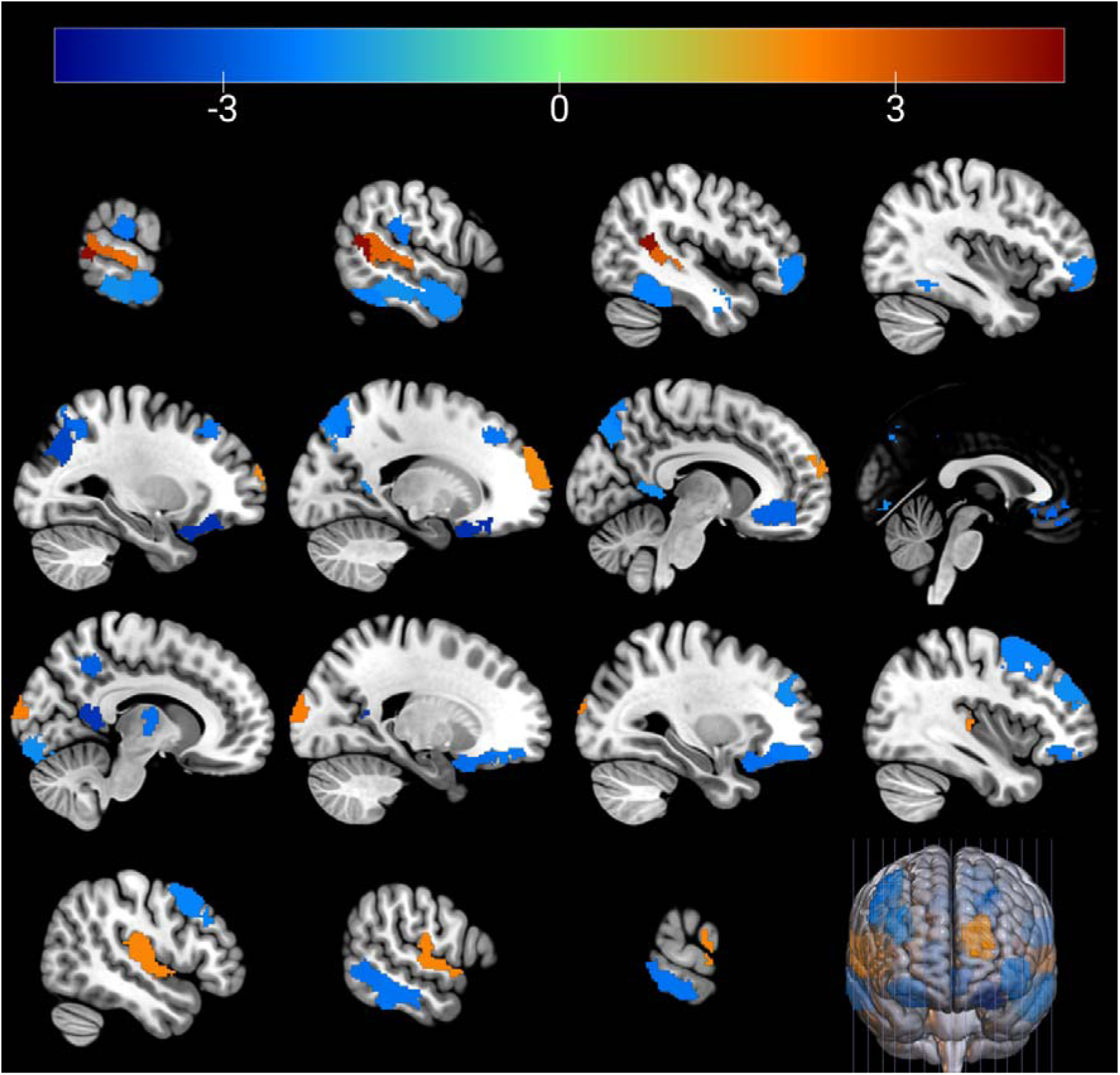
Group differences for modal controllability during 2-back task. ROIs are coloured by corresponding t-values using the same scale and colour conventions as Figure 6. Sagittal slice indices are as in Figure 6.

NF1 participants showed lower modal controllability than controls in the intraparietal sulcus, precuneus, lateral ventral and dorsal prefrontal cortex, and temporal region (control network); medial prefrontal cortex, dorsal prefrontal cortex, temporal cortex, and retrosplenial cortex (default mode network); superior parietal lobule and temporal occipital regions (dorsal attention network); orbitofrontal cortex (limbic network); parietal operculum and lateral prefrontal cortex (ventral attention network); extrastriate cortex (central visual network); and ventral posterolateral and medial dorsal thalamus (subcortical network). In contrast, NF1 participants showed higher modal controllability across secondary somatosensory and auditory cortices (somatomotor network), extrastriate cortex (central visual network), temporal parietal regions (temporal parietal network).

#### Activation energy

The region-specific t-statistics and p-values for activation energy are reported in full in Supplementary Table 5. Figure 8 presents the projection of these results to the brain template. NF1 participants showed lower activation energy than controls in the intraparietal sulcus, temporal region, precuneus, and lateral ventral prefrontal cortex (control network); medial prefrontal cortex, dorsal prefrontal cortex, temporal region and retrosplenial cortex (default mode network); superior parietal lobule and temporal occipital regions (dorsal attention network); temporal pole and orbitofrontal cortex (limbic network); and ventral posterolateral, intralaminar and medial dorsal thalamus and amygdala (subcortical network). In contrast, NF1 participants showed higher activation energy than controls in extrastriate cortex (central visual network), secondary somatosensory and auditory cortices (somatomotor network), temporal parietal regions (temporal parietal network), dorsal prefrontal cortex (default mode network), and anterior pulvinar thalamus (subcortical network).

**Figure 8.**
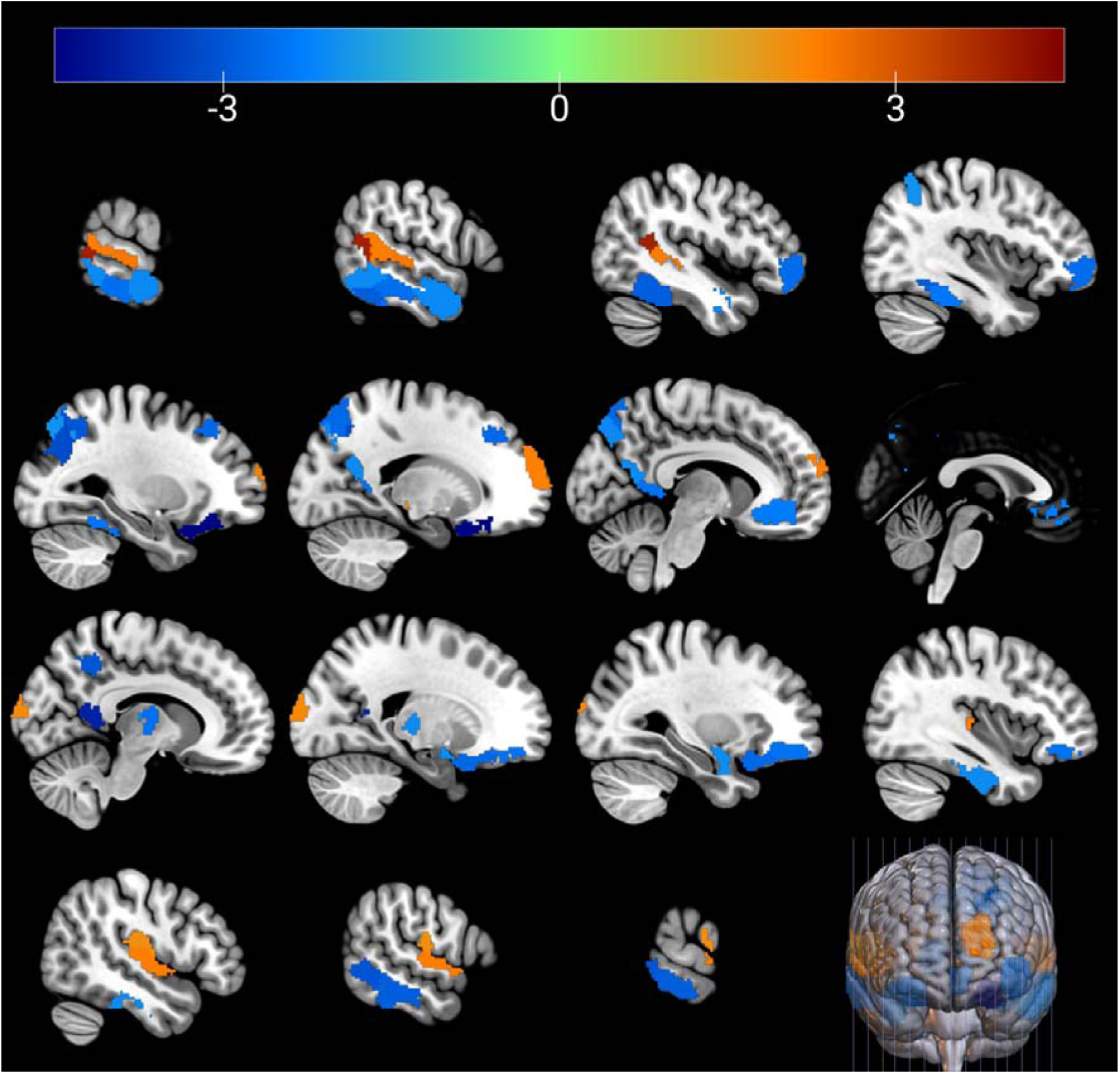
Group differences in activation energy during 2-back task. ROIs are coloured by corresponding t-values using the same scale and colour conventions as Figure 6. Sagittal slice indices are as in Figure 6.

### Region-wise controllability differences reliably classify individuals as NF1 or neurotypical controls

To determine whether the region-wise differences in controllability identified above were informative for distinguishing NF1 participants from neurotypical controls at the level of individual participants, we carried significant group differences in controllability measures forward to classification analysis.

All controllability measures showed the capacity to distinguish individual NF1 participants from neurotypical controls (AUC > 80%), with average controllability achieving the highest performance (76.9% balanced accuracy, 85.5% AUC). Combined models showed the greatest stability across repeated cross-validation (SD = 9.2% for balanced accuracy).

### Altered average controllability predicts individual differences in working memory performance in NF1

To test whether the region-wise controllability differences identified above were associated with individual differences in working memory performance, we used the significant regional group differences in controllability as predictors of a composite working-memory measure in the NF1 group. The composite was derived using PCA across the in-scanner and out-of-scanner working-memory tasks (Supplementary Material 3). Predictive performance was evaluated using 1,000 repetitions of cross-validation, with age and sex included as covariates.

Average controllability showed modest but statistically significant out-of-sample predictive performance, explaining 2.2% of variance in working-memory performance (R² = 0.022 ± 0.016; p = 0.049). Modal controllability, activation energy, and the combined model (including all three metrics) did not show significant out-of-sample predictive performance (Table 4, Supplementary Material 3). Thus, among the controllability measures that differed between groups, average controllability was the only measure that significantly predicted individual differences in working-memory performance in NF1.

**Table 4.**
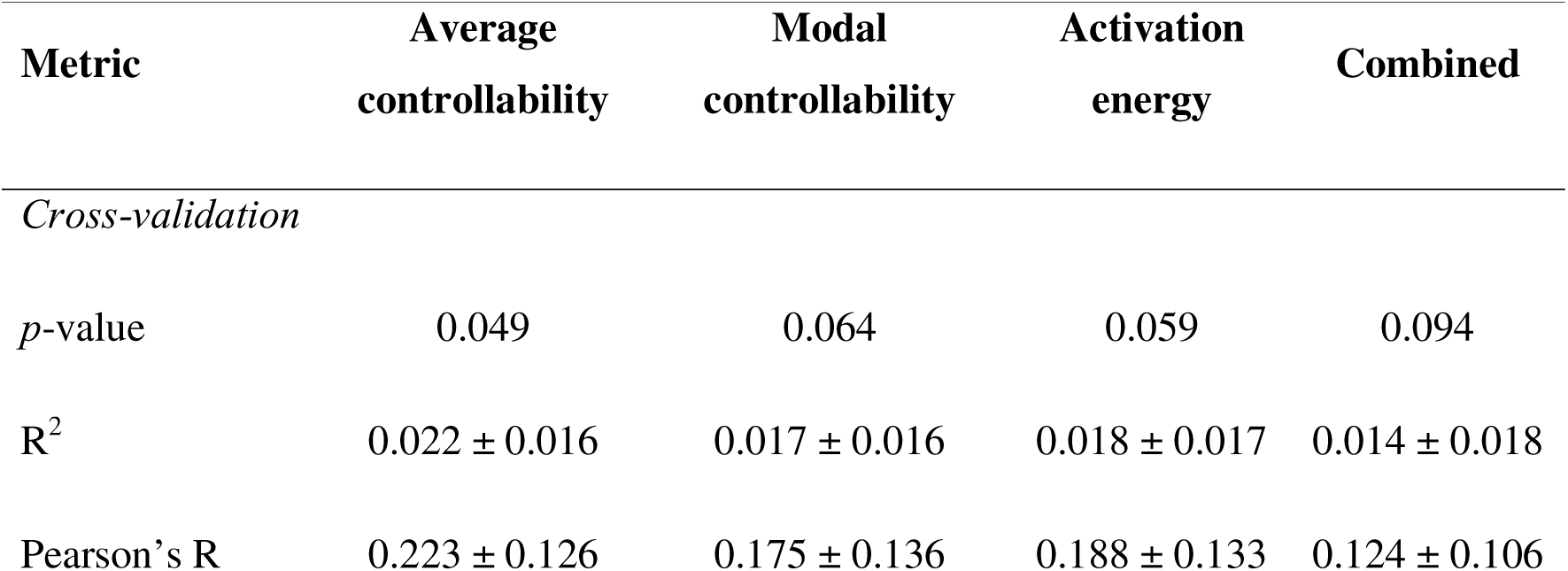
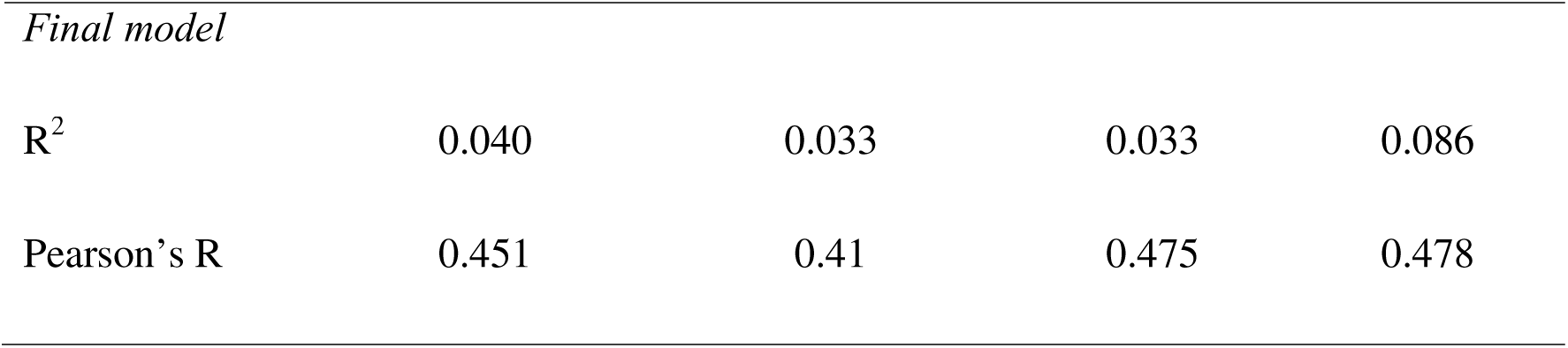
Behavioural prediction performance summary for models using network controllability measures. All models included age and sex as covariates. Values in cross-validation rows represent mean ± standard deviations across 1000 repetitions. Final models were fitted to the whole NF1 sample (n = 51). The p-values of predictive performance were estimated based on R^2^ obtained during permutation testing.

| **Metric** | **Average**<br>**controllability** | **Modal**<br>**controllability** | **Activation**<br>**energy** | **Combined** |
| --- | --- | --- | --- | --- |
| *Cross-validation* |  |  |  |  |
| *p*-value | 0.049 | 0.064 | 0.059 | 0.094 |
| $R^2$ | $0.022 \pm 0.016$ | $0.017 \pm 0.016$ | $0.018 \pm 0.017$ | $0.014 \pm 0.018$ |
| Pearson's R | $0.223 \pm 0.126$ | $0.175 \pm 0.136$ | $0.188 \pm 0.133$ | $0.124 \pm 0.106$ |
| *Final model* |  |  |  |  |
| R<sup>2</sup> | 0.040 | 0.033 | 0.033 | 0.086 |
| Pearson's R | 0.451 | 0.41 | 0.475 | 0.478 |

### Better working memory in NF1 is associated with lower average controllability in left prefrontal and superior parietal regions, and higher average controllability in right prefrontal, intraparietal, temporal-parietal, somatosensory and retrosplenial regions

The final models, fitted to the whole NF1 sample, were used to obtain interpretable regional beta weights. Figure 9 presents the beta weights for average controllability, while precise beta weights for all models are provided in Supplementary Material 3. Two predominant regional patterns were observed.

**Figure 9.**
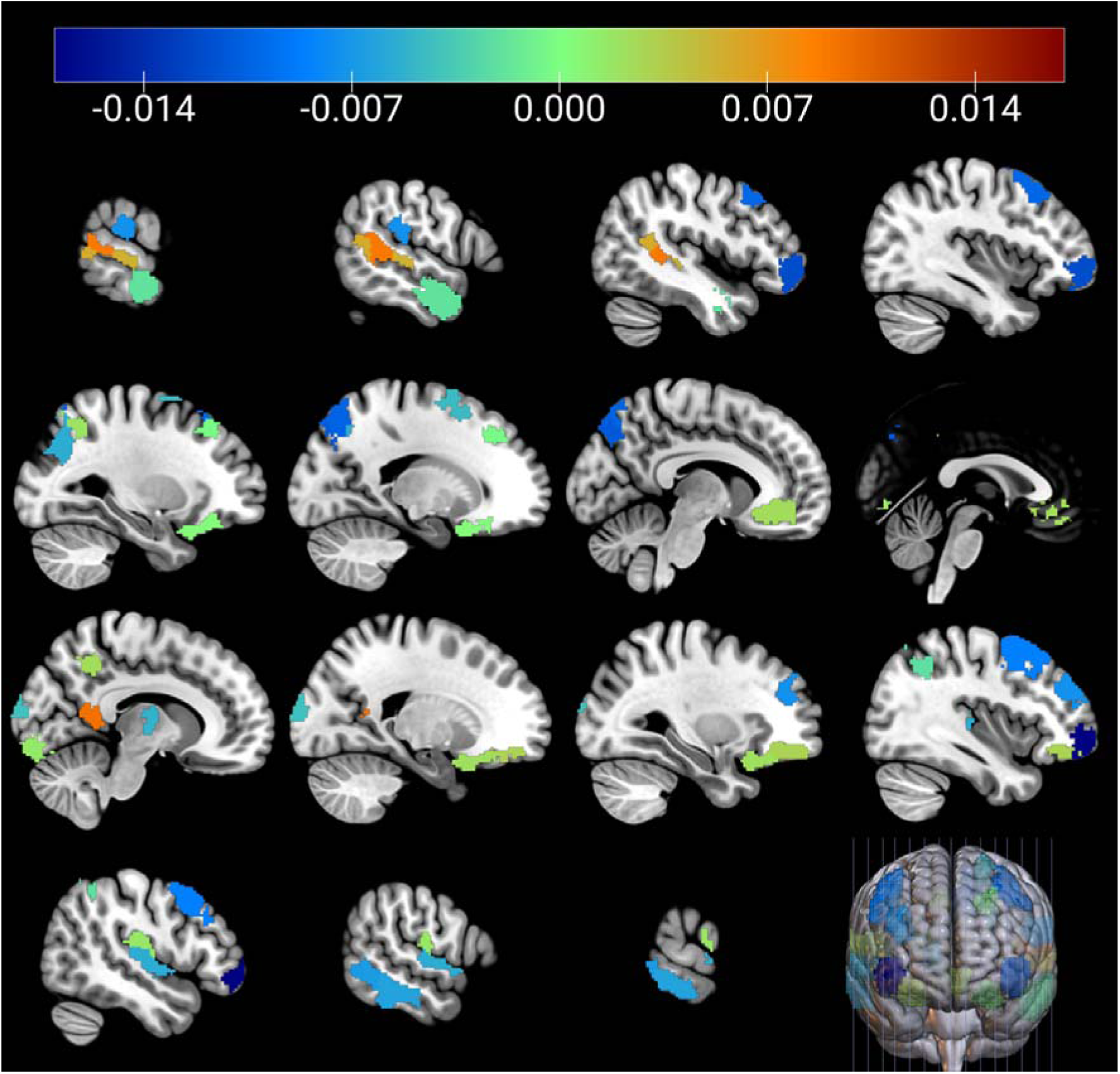
Average controllability beta weights from the final predictive model of the working memory component (N NF1 = 51). Weights have been projected onto brain surface for anatomical interpretability (colour range -0.017 to 0.017). Positive values indicate positive association between average controllability and working memory performance.

The first pattern showed that higher working memory performance was associated with lower average but higher modal controllability and activation energy in left lateral ventral prefrontal cortex (control B; ROI 104), left superior parietal lobule (dorsal attention A; ROIs 53 and 54), left temporal cortex (default B; ROI 123), and right ventral posterolateral thalamic nucleus (ROI 320). Additional regions showing this pattern but not reaching significant associations across all three measures included left precuneus (control C; ROI 108), right temporal region (control A and B; ROI 245 and 255) and right auditory cortex (somatomotor B; ROI 186).

The second pattern showed that higher working memory performance was associated with higher average but lower modal controllability and activation energy in left intraparietal sulcus (control A; ROI 92), right lateral ventral prefrontal cortex (control B; ROI 262), left temporoparietal cortex (temporal-parietal; ROIs 148 and 149), and right secondary somatosensory cortex (somatomotor B; ROI 192). Additional regions showing this pattern but not reaching significant associations across all three measures included right precuneus (control C; ROI 268), left temporoparietal cortex (temporal-parietal; ROI 150) and right retrosplenial cortex (default C; ROI 291).

Additional regions showing significant associations that did not conform to these two patterns included right extrastriate cortex (central visual; ROI 159), left medial and dorsal prefrontal cortex (default A; ROIs 112 and 119), right orbitofrontal cortex (limbic B; ROI 235), and left orbitofrontal cortex (limbic B; ROI 80).

## Discussion

This study is the first to characterise FC and network controllability during a working memory task in NF1 and a neurotypical control group, with the aim of using task-state network controllability to provide mechanistic insight into the network alterations underlying working memory impairments in NF1, and to establish whether these controllability profiles could serve as candidate markers for identifying neuromodulation targets. FC analysis revealed reduced connectivity within and between frontoparietal, dorsal attention and default mode networks in NF1, but increased connectivity between frontoparietal and somatomotor networks. Network controllability provided mechanistic insight into these findings, revealing how the regions of the network are positioned to transition the brain across states. We found that, compared to neurotypical controls, NF1 showed increased average controllability and reduced modal controllability and activation energy in regions within the control, default, and dorsal attention networks, particularly frontal and parietal regions. This hub-like profile typically facilitates transitions into common, easily reachable states and reflects regions that are more readily activated within, and able to spread activity across, the current network configuration, while reduced modal controllability indicates a reduced capacity to drive the brain towards the difficult-to-reach states required for coordinated activity during working memory performance (Deng et al., 2022; Gu et al., 2015). In contrast, sensorimotor and temporal-parietal networks showed lower average controllability, higher modal controllability and higher activation energy, indicating a more peripheral-like profile with greater capacity to facilitate specialised state transitions but less support from the current network configuration. Together, these findings suggest a redistribution of network control in NF1, in which canonical cognitive networks are biased towards readily accessible states while sensorimotor and temporal-parietal regions are better positioned to facilitate specialised transitions. Importantly, these altered control properties distinguished NF1 from controls, while average controllability predicted individual differences in working memory performance. These findings demonstrate that task-state network controllability is a behaviourally relevant and principled framework for identifying candidate neuromodulation targets.

### Network controllability is hierarchically organised in the neurotypical and NF1 brains

In neurotypical controls, average controllability increased from primary sensory and motor regions towards progressively higher-order association cortices. This pattern is similar to the hierarchical organisation of the human cortex, in which processing progresses from regions supporting relatively specific sensory representations towards higher-order association regions, integrating information across domains to support increasingly abstract, domain-general processes (Huntenburg et al., 2018; Margulies et al., 2016; Mesulam, 1998; Mesulam, 2008). This cortical pattern is also consistent with previous controllability research demonstrating that regions occupying more integrative, hub-like positions within the network tend to exhibit greater average controllability and facilitate transitions towards easily reachable states (Gu et al., 2015). Subcortical structures showed some of the highest average controllability. Modal controllability showed the opposing spatial pattern, with higher values in sensory-motor regions and lower values in higher-order association cortex and subcortical structures. This pattern is consistent with studies showing that relatively peripheral regions are better positioned to facilitate transitions towards difficult-to-reach states (Deng et al., 2022; Gu et al., 2015). Together, these findings indicate that neurotypical functional networks show distributed control capacity across a cortical hierarchy, with higher-order association regions supporting easy-to-reach configurations and lower-order regions having greater capacity for specialised state transitions.

Activation energy demonstrated a unique spatial pattern, characterised by relatively lower values in subcortical structures and tendency towards higher values across cortical association networks. This distribution suggests that higher-order association cortices may require more energy from the network to transition across states, consistent with their role in integrating information across distributed systems and supporting flexible, context-dependent cognitive processes (Huntenburg et al., 2018; Margulies et al., 2016; Mesulam, 1998; Mesulam, 2008). Conversely, the lower energy observed in subcortical and sensory regions may reflect their embedding within more readily accessible network configurations and their contribution to maintaining ongoing patterns of brain dynamics (Deng et al., 2022; Saberi et al., 2024).

### Frontoparietal regions are disconnected and biased toward routine states in NF1

Both FC and controllability demonstrated a substantial disruption in the function of the frontoparietal network in NF1 during a verbal working memory task. NF1 participants showed reduced connectivity, as compared to controls, between left prefrontal cortex and left intraparietal sulcus and bilateral inferior parietal lobule, which extends previous reports of reduced activation (hypoactivation) in NF1 groups in these regions during working memory (Ibrahim et al., 2017; Shilyansky et al., 2010). These regions are usually involved with goal maintenance, retrieval, and flexible attentional shifting during working memory (Owen et al., 2005), which suggests that the canonical working memory network does not function as a cohesive, integrated circuit in people with NF1. There were further disruptions to connectivity between this lateral prefrontal cortex of the frontoparietal network and left superior parietal lobule of the dorsal attention network. Typically these regions support top-down goal directed attention during cognitive tasks (Corbetta & Shulman, 2002), which further suggests that people with NF1 have compromised connectivity between executive control and goal-directed attention systems. Together our findings of FC disruptions extend the previous literature in NF1 by demonstrating substantial loss of connectivity during working memory between network regions supporting executive control, goal-directed behaviour and top-down attentional selection. Additionally, the disruptions of left prefrontal cortex connectivity are also consistent with reports of reduced anterior-posterior long-range connectivity in NF1 (Baudou et al., 2020; Mennigen et al., 2019), though it is unknown whether this reflects specific vulnerability of left prefrontal cortex (Litwińczuk et al., 2026).

Network controllability analysis was performed to investigate how effectively brain regions drive transitions between neural states. This provides a mechanistic link between regional connectivity properties and their functional role in supporting cognitive transitions. Importantly, we must highlight that because these controllability profiles were estimated from functional connectivity measured during the 2-back condition, they reflect the organisation of the functional network while participants were actively performing the working memory task, rather than intrinsic resting-state or structural architecture. Such characterisation is relatively novel but has been successfully performed in other reports (Wang et al., 2025; J. Yang et al., 2025; Y. Yang et al., 2025), providing unique insights about task-dependent functional organisation in neurotypical and clinical populations.

Our analysis revealed that in the NF1 group, relative to controls, frontoparietal regions show higher average controllability, lower modal controllability, and lower activation energy. This means that the frontoparietal regions, which are typically well suited to drive the network towards demanding states due high modal controllability (Gu et al., 2015), were biased towards hub-like dynamics in NF1. Lower activation energy in these regions reflects that they are readily activated within the current network organisation. Together, these patterns of controllability differences suggest that in NF1, these regions are positioned in the network in such way that are easily engaged in low-energy, routine brain states, but cannot effectively steer the brain toward specialised, difficult-to-reach states.

In our previous work, we showed that NF1 participants have increased endogenous self-connectivity within the left dorsolateral prefrontal cortex and inferior parietal gyrus, reflecting reduced sensitivity of these regions to external inputs during both high and low working memory loads (Litwińczuk et al., 2026). The present controllability results further suggest that this reduced responsiveness may be related to a broader functional network connectivity alteration in NF1. Specifically, the network topology is altered such that these frontoparietal regions are integrated into a dense core that favours routine dynamics, rather than being positioned as peripheral drivers of specialised states. Future research combining neurotransmitter activity and network controllability measures could clarify whether the hub-like dynamics identified in this work reflect the GABAergic dysregulation associated with NF1 (Violante et al., 2012). Mechanistically, an altered inhibitory landscape might widen the energy gap between easy- and difficult-to-reach states, thereby increasing the metabolic cost of demanding cognitive transitions and rendering the network more rigid.

Overall, the FC and controllability patterns of the frontoparietal network in NF1 suggest that the network is not only hypoactive but also disconnected and organised in a way that reduces the capacity of these regions to steer the network toward demanding cognitive states.

### Altered connectivity and network control profiles explain reduced suppression of default mode network in NF1

This study also provides further evidence for the default interference hypothesis (Violante et al., 2012), which proposes that the default mode network is insufficiently suppressed during performance in NF1, thereby interfering with the activity of task-relevant networks. Previous work showed that increased activation of the PCC and temporal regions, together with increased PCC-visual network connectivity, were associated with visuospatial working memory performance in NF1 (Ibrahim et al., 2017). Here, FC analysis showed that the default mode network was internally fragmented, with reduced connectivity between left dorsal prefrontal and left parahippocampal cortices, and between the left parahippocampal and right temporal cortices. In addition, the default network showed increased connectivity between the medial prefrontal cortex and frontal regions of the frontoparietal network.

Together, the fragmented internal connectivity and increased connectivity with frontoparietal regions suggest that the default mode network in NF1 is reorganised in a way that may increase the accessibility of default mode regions during working memory, rather than reflecting a coherent shift towards rest-like activity. Network controllability analysis allowed us to test this by quantifying the functional consequences of this altered connectivity for the network’s capacity to transition between states during the task.

Network controllability analysis of the default mode network revealed a pattern similar to that observed in the frontoparietal network, characterised by higher average controllability, lower modal controllability and lower activation energy in medial prefrontal, dorsal prefrontal, temporal and retrosplenial regions. Deng et al. (2022) and Saberi et al. (2024) showed that, in neurotypical individuals, network topology during task performance increases the energetic cost of activating task-irrelevant regions, reflected by higher activation energy in default mode regions. In contrast, the lower activation energy observed in the NF1 group suggests that the altered connectivity identified by the FC analysis leads to network failure to increase the energetic cost of activating the default mode network regions during working memory. This finding is consistent with impaired suppression of the default mode network during task performance, as proposed by the default mode interference hypothesis (Violante et al., 2012).

Our findings provide a functional account of the default mode interference hypothesis, suggesting that during working memory the default mode network regions in NF1 are atypically connected to the rest of the network, reflecting the fragmented intra-DMN connectivity and increased coupling with frontoparietal regions described above. Due to altered FC patterns in NF1, these regions are more readily activated within the existing network architecture and better positioned to steer the brain toward a broad repertoire of low-energy, routine states during working memory, consistent with their higher average controllability relative to neurotypical controls. This framework provides a mechanistic interpretation of previous observations of reduced DMN suppression in NF1 (Ibrahim et al., 2017), while further supporting the default mode interference hypothesis (Violante et al., 2012).

Importantly, one caveat must be considered that because network controllability was estimated from functional connectivity measured during the 2-back condition, these control properties reflect the organisation of the functional network while participants were actively performing the working memory task, rather than intrinsic resting-state architecture. Comparison of the resting-state and task-specific controllability patterns in NF1 will further help determine whether these alterations represent stable features of network organisation or whether they are specifically related to cognitive demands.

### Sensorimotor and temporal parietal regions are biased toward specialised, high-cost states in NF1

The FC analysis revealed that the NF1 group exhibited increased connectivity between right somatomotor cortex and left lateral prefrontal cortex, left postcentral cortex, and left insula, together with increased connectivity between temporal parietal and somatomotor networks at the network level. These task-state alterations raise the question of whether this network reorganisation reflect a compensatory mechanism that supports working memory performance in NF1, or, alternatively, maladaptive connectivity that interferes with task processing, analogous to the default mode interreference hypothesis discussed above.

Network controllability analysis further showed that the NF1 group exhibited increased modal controllability and activation energy in secondary somatosensory and auditory cortices, the central visual extrastriate cortex, and regions of the temporal parietal network. These findings suggest that, during working memory, these regions exhibit control properties more characteristic of peripheral nodes, with an increased capacity to drive the network towards specialised, difficult-to-reach states, at a higher energetic cost (Gu et al., 2015). The increased FC between somatomotor and frontoparietal regions with this redistribution of control properties, suggesting that during working memory sensorimotor regions of individuals with NF1 become better positioned to drive specialised state transitions, while the frontoparietal regions have characteristics more typical of network hubs. During working memory, this organisation creates a mismatch between regions that are theoretically better positioned to facilitate demanding state transitions and regions that remain biased towards routine, readily accessible network configurations. Interestingly, this pattern contrasts with recent network controllability findings in schizophrenia (J. Yang et al., 2025), where both frontoparietal and sensorimotor systems showed a generalised reduction in controllability. This contrast suggests that the altered control properties of sensorimotor regions in NF1 are unlikely to simply reflect a non-specific consequence of cognitive impairment or globally reduced controllability. Rather, they are consistent with a task-specific reorganisation of network control during working memory. Although NF1 and schizophrenia both exhibit working memory deficits and frontoparietal abnormalities, these contrasting controllability profiles suggest that the network mechanisms underlying these impairments may differ substantially.

Taken together, the FC and controllability findings suggest that during working memory the sensorimotor and temporoparietal systems in NF1 adopt a control profile that differs substantially from that of neurotypical controls. Increased modal controllability and activation energy suggest that these regions become better positioned to facilitate specialised state transitions within the task-state network, while frontoparietal regions retain a more hub-like organisation. Whether this redistribution of control supports compensation or instead contributes to inefficient task processing remains unclear. However, the findings clearly indicate that working memory in NF1 is associated with a distinct reorganisation of functional network control.

#### Classification and prediction

An important implication of these findings is that the altered network control properties identified in NF1 are sufficiently robust to generalise to individual participants. Classification analyses demonstrated that average controllability, modal controllability and activation energy distinguished participants with NF1 from controls, while prediction analyses showed that average controllability was also associated with individual differences in working memory.

These findings demonstrate that differences in average controllability, modal controllability and activation energy are sufficiently consistent across individuals to distinguish participants with NF1 from controls, with all classifiers achieving good discriminative performance (AUC > 80%) and average controllability emerging as the strongest individual predictor. Sensitivity consistently exceeded specificity, indicating that the controllability measures more reliably identified participants with NF1 than neurotypical controls. This pattern suggests greater heterogeneity among controls, some of whom exhibited controllability profiles overlapping with those observed in NF1. Adjusting for age and sex had minimal impact on classification performance, indicating that these findings were not driven by demographic differences. The combined model achieved the highest accuracy and lowest cross-validation variability, indicating that the three controllability measures complementary rather than redundant information.

Average controllability within regions showing altered network control in NF1 also significantly predicted working memory performance in unseen participants, whereas modal controllability and activation energy showed trends towards significance. Importantly, the relationship between average controllability and cognition was region-dependent rather than uniform. Greater average controllability within canonical cognitive control regions (Owen et al., 2005) predicted poorer working memory, whereas higher average controllability in selected default mode and limbic regions predicted better performance. This suggests that the behavioural consequences of altered controllability depend on the functional role of the affected regions rather than reflecting a global increase or decrease in network control.

Specifically, higher average controllability in left lateral ventral prefrontal cortex and left superior parietal lobule predicted poorer working memory performance. These were the same regions showing increased average controllability in NF1, consistent with a shift away from the strong control properties these regions typically exhibit in neurotypical individuals (Gu et al., 2015). In contrast, greater average controllability in right lateral ventral prefrontal cortex and left intraparietal sulcus predicted better performance, raising the possibility that increased hub-like dynamics in these regions reflects compensatory network reorganisation, though whether this represents NF1-specific compensation or beneficial variation independent of the disorder remains unclear.

This region-dependent pattern indicates that the behavioural outcomes of altered controllability in NF1 depend on the functional role of the affected regions, rather than reflecting a uniform increase or decrease in network control. Together with the classification results, these findings position average controllability not merely as a discriminative marker of NF1, but as a measure that captures functionally meaningful, region-specific network alterations linked to working memory performance, identifying which network alterations are behaviourally relevant and therefore warrant further mechanistic investigation.

#### Directions for future research

A key implication of these findings is that task-state network controllability provides a principled framework for identifying candidate neuromodulation targets in NF1. Unlike conventional activation and FC analyses, controllability metrics quantify how regional interactions influence the brain’s ability to transition between functional states, thereby offering mechanistic insight into why particular network alterations are associated with impaired working memory. Our findings provide a coherent picture of disrupted FC and network controllability in NF1. In particular, left lateral ventral prefrontal cortex, right lateral dorsal prefrontal cortex, and left intraparietal sulcus consistently exhibited a more hub-like control profile (i.e. higher average controllability, lower modal controllability, and lower activation energy), and the magnitude of these alterations predicted poorer working memory performance. These findings raise the possibility that interventions capable of modifying the control properties of specifically these regions may improve working memory performance in NF1.

Importantly, however, controllability metrics describe properties of the network as a whole, rather than isolated regional characteristics. Consequently, the altered control profiles identified here are likely to arise from distributed changes in functional connectivity rather than local cortical dysfunction alone. An important next step will be to identify which specific connectivity changes drive the altered controllability profiles observed in NF1. For example, a region’s disrupted controllability may reflect local dysfunction and/or changes in other parts of the network, such as an altered connection or a change in the excitability of another node, that render the network to be more resistant to being controlled by that region. Establishing this relationship will also be important for interpreting the effects of non-invasive brain stimulation. Combining stimulation with network controllability analyses could determine how different stimulation protocols reshape task-state functional networks and whether these changes translate into improvements in cognitive performance (Garg et al., 2022).

In this study, network controllability was only assessed during the high-load (2-back) condition. Future studies should characterise how network controllability is reconfigured across increasing levels of cognitive demand to determine whether NF1 is characterised by impaired transitions between control states rather than abnormalities confined to high task load. Such paradigms would also allow testing whether brain stimulation facilitates the dynamic reconfiguration of network control across cognitive demands, rather than simply altering the control properties of individual regions within a single task condition. Finally, it will be important to determine how the altered control properties of frontoparietal regions interact with the default mode and limbic networks, whose increased average controllability was associated with better working memory performance and may therefore reflect compensatory network reorganisation. Therapeutic strategies that modulate frontoparietal controllability without accounting for these potentially compensatory mechanisms may prove ineffective or even impair cognitive performance.

### Strengths and limitations

Despite the challenges associated with a relatively small paediatric clinical cohort, the study produced robust classification and predictive models based on network controllability measures. The NF1 cohort was characterised by relatively high in-scanner motion, heterogeneous medication use, and frequent co-occurrence of attention deficit hyperactivity disorder (ADHD) and autism spectrum conditions. Nevertheless, including mean framewise-displacement, diagnostic comorbidities and medication categories (antidepressants, antipsychotics, stimulants) as covariates had minimal on model performance, indicating that predictive value of the controllability measures was robust to these sources of heterogeneity within this cohort.

A further strength is that, despite ceiling effects in behavioural performance during the in-scanner verbal 2-back task, network controllability measured during the 2-back condition predicted a broader working memory construct incorporating in-scanner performance (0-back and 2-back), visuospatial working memory, and memory span. This suggest that the identified controllability alterations generalise beyond the specific task performed during scanning and relate to broader working memory ability.

Several limitations should also be acknowledged. First, controllability was estimated only from the 2-back condition, providing a snapshot of task-state network organisation. Consequently, the present study cannot determine whether the observed alterations are specific to high working memory load, extend across cognitive tasks, or reflect more general properties of functional brain organisation in NF1. In addition, developmental trajectories may vary according to NF1 mutation type, disease severity, and medication exposure. In particular, stimulant medication may influence dopaminergic and noradrenergic systems known to modulate FC (Dipasquale et al., 2020), potentially affecting controllability estimates.

From a methodological perspective, controllability was derived from undirected, symmetric FC, which does not capture the directional asymmetry of neural interactions. In addition, the analyses relied on a linear network control framework, whereas brain dynamics are inherently nonlinear. Although previous work suggests that linear approximations capture meaningful aspects of large-scale brain dynamics, controllability estimates remain dependent on this modelling assumption (Tu et al., 2018). Finally, the concept of activation energy in network control theory should not be interpreted as a direct measure of neural or metabolic energy expenditure. Rather, it represents a mathematical quantity describing the relative effort required to drive transitions between network states within the model (Tu et al., 2018). Accordingly, the controllability metrics reported here should be interpreted as relative properties of task-state network organisation rather than absolute measures of the energetic demands of working memory. The present findings therefore characterise differences in network control between NF1 and neurotypical sample, rather than absolute physiological energy requirements.

Finally, although network controllability was originally developed for structural brain networks, the present study applied the framework to task-state functional connectivity, replicating the approach of several recent studies (Wang et al., 2025; J. Yang et al., 2025; Y. Yang et al., 2025). This approach provides insight into the control properties of the functional network supporting working memory, but the resulting controllability measures should be interpreted as properties of the observed functional architecture rather than of the underlying anatomical connectome.

## Conclusions

This study demonstrates that working memory in NF1 is characterised by a systematic reorganisation of task-state functional network control. This reorganisation was characterised by disrupted connectivity within and between the frontoparietal, dorsal attention, and default mode networks, accompanied by altered controllability profiles that favoured readily accessible network states over demanding state transitions. Conversely, sensorimotor and temporoparietal systems exhibited increased connectivity and greater capacity to facilitate demanding state transitions, albeit at higher theoretical energetic cost. Together, these findings suggest a redistribution of control across large-scale brain networks, creating a functional mismatch in which canonical cognitive control systems become biased towards routine network configurations, whereas sensorimotor systems assume greater capacity to facilitate demanding state transitions. Importantly, these altered control properties reliably distinguished individuals with NF1 from neurotypical controls, and average controllability predicted individual differences in working memory performance. Collectively, these findings identify altered task-state network controllability as both a mechanistic marker of cognitive dysfunction and a predictor of working memory performance in NF1, providing a foundation for future studies investigating targeted neuromodulation and other interventions aimed at restoring functional network control.

By combining task-state functional connectivity with network controllability analysis, this study moves beyond describing where network abnormalities occur to characterising how they alter the brain’s capacity to transition between functional states during working memory. In doing so, it provides a mechanistic framework linking large-scale network organisation to cognitive dysfunction in NF1 and establishes network controllability as a promising tool for identifying biomarkers and evaluating future therapeutic interventions.

## Methods

### Participants

The data for NF1 participants includes and expands on the data from a previous study described in detail in Garg et al. (2022) resulting in a total sample size of 53. All recruitment of NF1 participants was done via the Northern UK NF- National Institute of Health, with (i) diagnostic criteria [National Institutes of Health Consensus Development Conference. Neurofibromatosis conference statement. Arch. Neurol. 45, 575–578 (1988).] and/or molecular diagnosis of NF1; (ii) no history of intracranial pathology other than asymptomatic optic pathway or other asymptomatic and untreated NF1-associated white matter lesion or glioma; (iii) no history of epilepsy or any major mental illness; and (iv) no MRI contraindications. Participants on pre-existing medications such as stimulants, melatonin or selective serotonin re-uptake inhibitors were not excluded from participation. Additionally, 36 adolescents with no diagnosis of NF1 were recruited from staff newsletter and via local schools. The inclusion criterion for the control group was no history of neurodevelopmental conditions. Controls’ age and sex were matched to the sample of NF1 participants. Table 1 includes demographics information for NF1 participants and neurotypical controls.

### Working memory task

The verbal N-back task was used to assess working memory performance in the participants (Kirchner, 1958) (Figure 10). Within the MRI scanner, participants were presented with a sequence of black coloured letters on a grey screen. The participants were instructed to respond only to the target by pressing a handheld button. During the 0-back condition, the participants responded when the letter ‘X’ was presented on the screen. During the 2-back condition, the participants responded when letter on the screen matched the letter presented 2 screens before. The stimuli were presented for 500 milliseconds (ms) and there was a 2500 ms inter-stimulus interval. Each session consisted of 6 blocks of 0-back condition interleaved with 6 blocks of 2-back condition, each block lasted for 30 seconds and consisted of 9 trials. Before each block, an instruction screen displayed “0-back” or “2-back” for 2000ms to inform the participants about the upcoming condition. Participants completed a practice session before entering the scanner, which was monitored by a researcher to ensure that the participants understood the task correctly.

**Figure 10.**
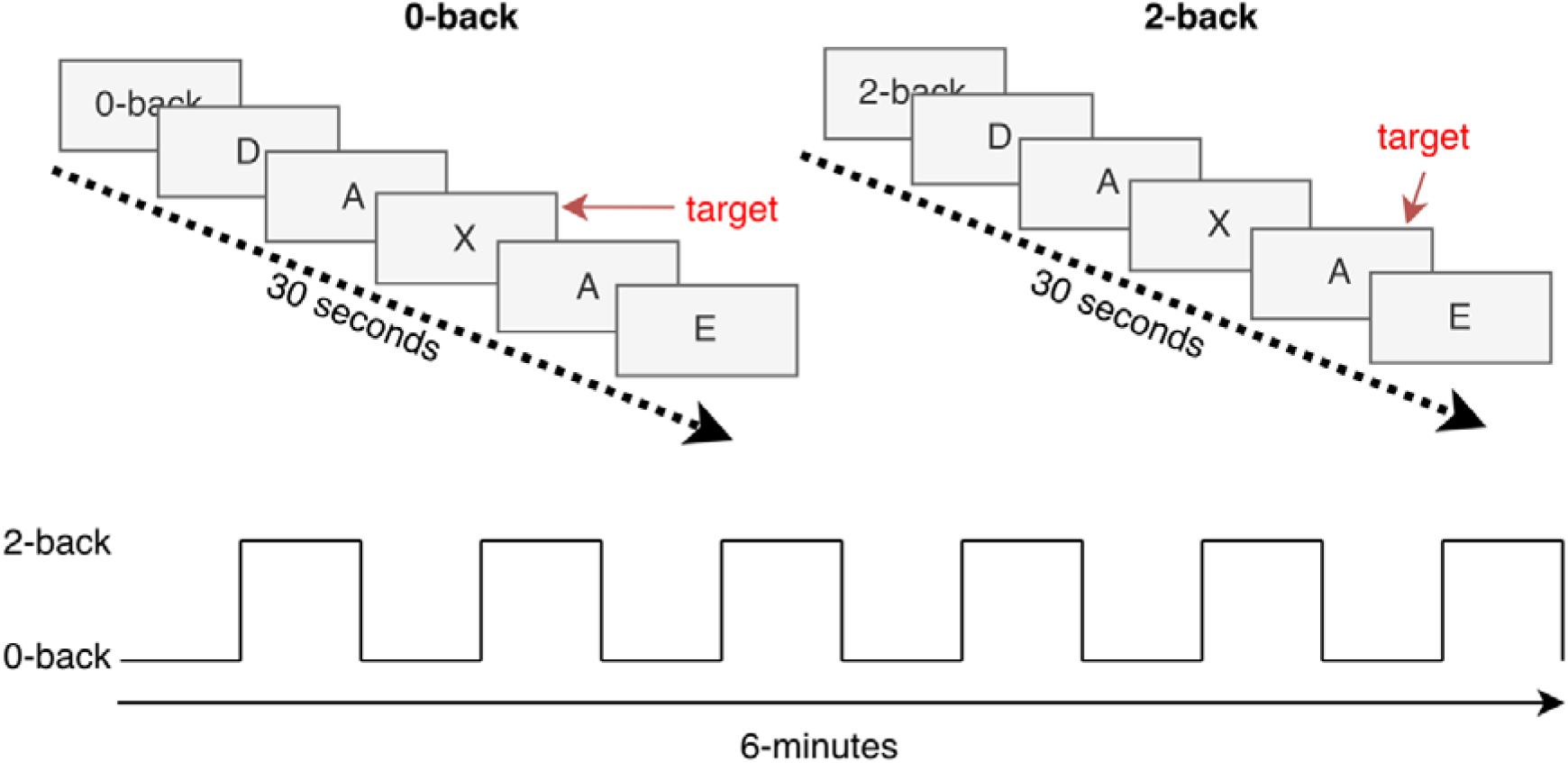
A schematic illustration of the experimental task design.

All measures of behavioural performance were calculated separately for 0-back and 2-back conditions. Accuracy was calculated as (correct hits + correct rejections)/total trials. D-prime was calculated as a measure of perceptual sensitivity, representing the ability to discriminate between target and non-target stimuli while accounting for response bias. It was calculated as z(hit rate) - z(false alarm rate), where z represents the inverse of the standard normal cumulative distribution function (Macmillan & Creelman, 2004). Response times (RT) were calculated only for correct responses to target stimuli. Inverse efficiency score (IES) was calculated by dividing RT by accuracy as a measure of speed-accuracy trade-off, in which lower scores reflect better cognitive performance (faster response for less accuracy cost) (Bruyer & Brysbaert, 2011). The verbal N-back data for 10 control and 2 NF1 participants were unavailable.

### Additional behavioural measures

Prior to scanning, in a quiet room, the participants completed two additional behavioural tasks: visuospatial N-back task and Corsi Blocks task. Out-of-scanner behavioural measures were not available for 2 NF1 participants.

#### Visuospatial N-back task

This task was presented using E-Prime 3.0 based on Jaeggi et al. (2010). On a computer screen, the participants were presented sequentially with a blue square in one corner of a 2x2 grid. A fixation cross remained present at the centre of the grid through the task. Stimulus appeared for 500ms, followed by a 2,500ms inter-stimulus interval (total trial duration: 3,000ms). The task progressed in difficulty from 0-back to 3-back. During 0-back condition, the participants responded when the yellow square appeared. For all other conditions, the square remained blue through all the trials and the participants responded whenever square location matched that presented N-trials before (where N = 1, 2, 3). The participants made their response using the ‘a’ key on the keyboard (marked with a tape).

Before each block, the participants were presented with written instructions on the screen (Figure 11) and had a chance to practice before the first experimental block started. Each block started with 2 start items that were never target trials, followed by 20 trials, of which 8 were targets. There were 3 blocks for each condition, presented in ascending difficulty level, presenting 240 trials in total. After each block the participants were informed of their accuracy and had a moment to rest before progressing. All four measures of behavioural performance (accuracy, response time, D-prime and IES) were calculated for this task in the same way as the verbal N-back task.

**Figure 11.**
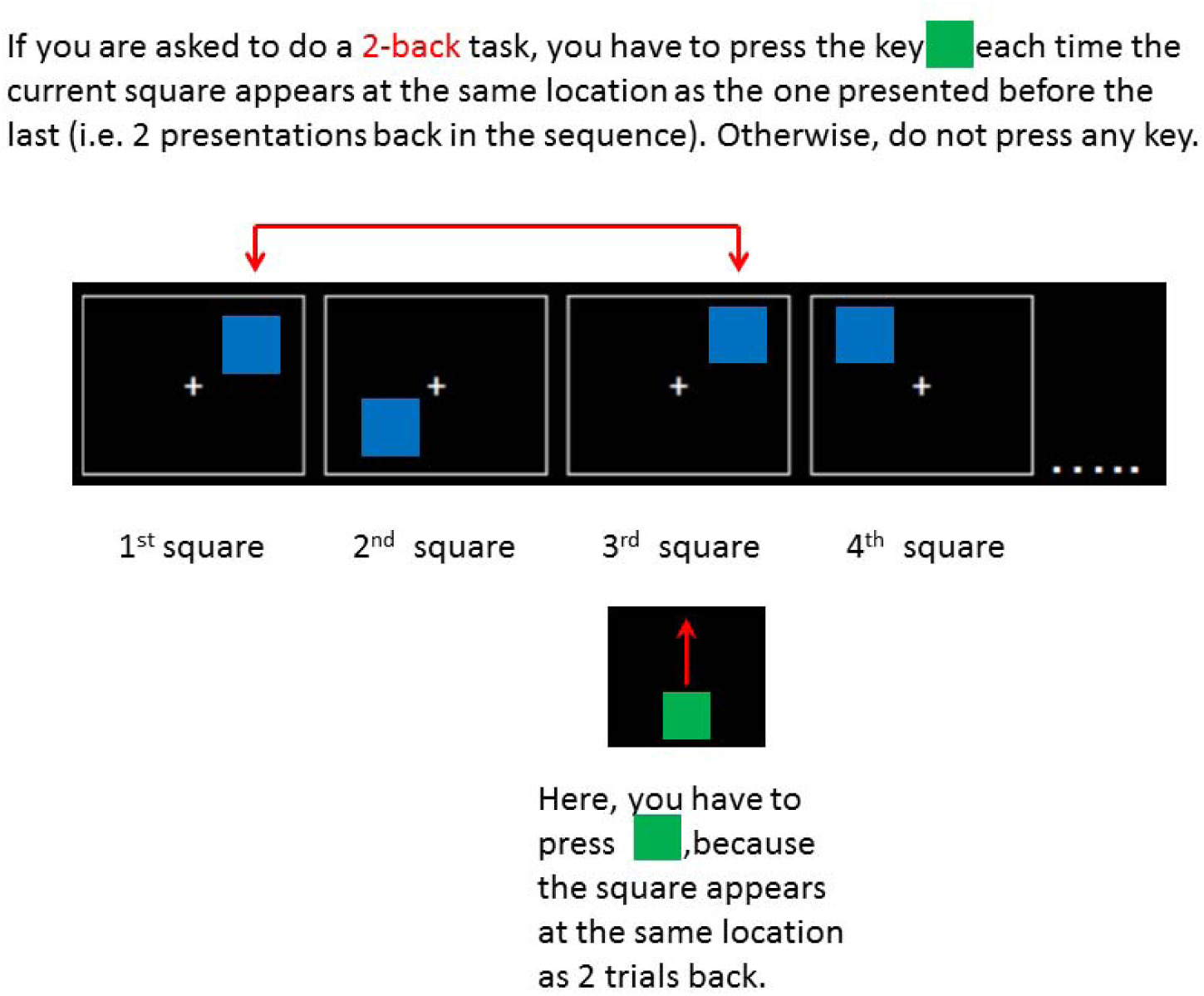
The instruction screen from the 2-back condition of the visuospatial N-back task.

#### Corsi Blocks

This task was presented using the Psychology Experiment Building Language (PEBL 2.1). During the Corsi Blocks task the participants were presented with 9 blue squares on a black background. The squares sequentially changed colour to yellow for 1000ms, the time between squares flashing in yellow was 1000ms and inter-trial interval 1000ms. The participants were asked to remember the sequence in which the squares were highlighted and reproduce it using a mouse button, followed by pressing “Done” at the bottom of the screen to indicate they completed their response. The cursor was hidden during sequence presentation, and it showed and repositioned to screen centre at start of each response. The sequence length started with 2 colour changes and increased up to 9; the length increased if participants correctly reproduced at least one of the two trials at a given length. The participants completed 3 practice trials, with sequences of 3 flashes. Participant performance was measured as memory span, which is a measure of average recall performance across the task and is calculated by PEBL as (2 + number of correct trials)/2. Two NF1participants had Corsi block memory span resulting with 51 datapoints, and one neurotypical control participant’s memory span was not available resulting with 35 remaining datapoints.

### MRI acquisition

The data collected here is part of a longer MRI acquisition session, registered at https://clinicaltrials.gov/ct2/show/NCT04991428. The complete acquisition sequence is presented Figure 12.

**Figure 12.**
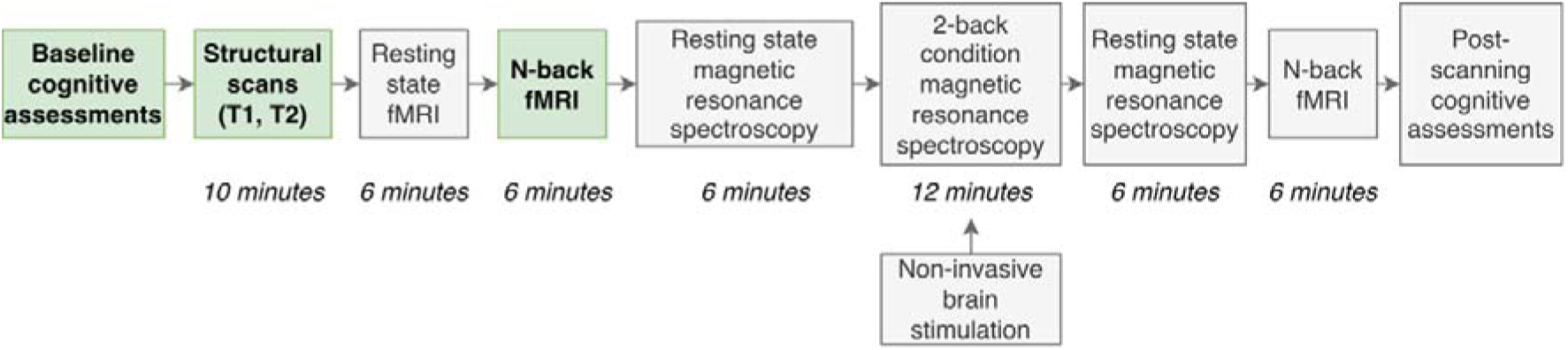
The complete MRI acquisition sequence. Green squares were used to indicate the data used as part of this report.

Structural scanning was conducted using a Philips Achieva 3 T MRI scanner (Best, NL) equipped with a 32-channel head coil. First, 3D T1-weighted magnetic resonance images were obtained in the sagittal plane with a magnetization-prepared rapid acquisition gradient-echo sequence (repetition time = 8.4 ms; echo time = 3.77 ms; flip angle = 8°; inversion time = 1150 ms; in-plane resolution = 0.94 mm; 150 slices with 1 mm thickness). Next, a T2-weighted structural scan was performed using a turbo spin echo sequence (TR = 3756 ms; TE = 89 ms; 40 slices of 3 mm thickness and 1 mm gap; in-plane resolution = 0.45 mm).

Functional imaging was performed on a 3 Tesla Philips Achieva scanner using a 32-channel head coil with a SENSE factor 2.5. To maximise signal-to-noise (SNR), we utilised a dual-echo fMRI protocol developed by Halai et al. (2014). The fMRI sequence included 36 slices, 64×64 matrix, field of view (FOV) 224×126×224 mm, in-plane resolution 2.5×2.5 mm, slice thickness 3.5 mm, TR=2.5 s, TE = 12 ms and 35 ms. The total number of volumes collected for each fMRI session was 144.

### MRI Processing

Image processing was done using SPM12 (Wellcome Department of Imaging Neuroscience, London; http://www.fil.ion.ucl.ac.uk/spm) and MATLAB R2023a. Dual echo images were extracted and averaged using in-house MATLAB code developed by Halai et al. (2014) (DEToolbox). First, functional images were slice-time corrected and realigned to first image. Then, the short and long echo times were averaged for each timepoint. The orientation and location of origin point of every anatomical T1 image was checked and corrected where needed. First functional EPI image was co-registered to the structural (T1) image. Motion parameters estimated during co-registration of long echo-time images were input to Artifact Detection Tools (ART; https://www.nitrc.org/projects/artifact_detect/) toolbox along with combined dual echo scans for identification of outlier and motion corrupted images across the complete scan. The outlier detection threshold was set to changes in global signal 3 z-scores away from mean global brain activation. Motion threshold for identifying scans to be censored was set to 3 mm. Outlier images and images corrupted by motion were censored during the analysis by using the outlier volume regressors. Participants with less than 80% of scans remaining were removed from analysis. Following the removal of participants with high motion and acquisition artefacts, 53 NF1 and 36 neurotypical participants remained.

Unified segmentation was conducted to identify grey matter, white matter, and cerebrospinal fluid. Normalisation to MNI space was done with diffeomorphic anatomical registration using exponentiated lie-algebra (DARTEL) (Ashburner, 2007) registration method for fMRI. Normalised images were interpolated to isotropic 2 × 2 × 2 mm voxel resolution. A 6x6x6mm full width at half maximum (FWHM) Gaussian smoothing kernel was applied.

Functional denoising was performed using the Conn toolbox (Whitfield-Gabrieli & Nieto-Castanon, 2012), we followed the default fMRI denoising pipeline (Nieto-Castanon, 2020). First, aCompCor removed confounding effects of signal from white matter and cerebrospinal areas, session and task effects, and subject-motion parameters (3 translation and 3 rotation parameters) and global signal outlier scans, and session and task effects (Behzadi et al., 2007). Next, the denoising pipeline performed band-pass filtering (0.009–0.08 Hz) (Power et al., 2012; Yamashita et al., 2018).

Functional connectivity was estimated for 354 regions of interest (ROIs) using 300 cortical parcels from Schaefer et al. (2018) (Schaefer-300) and 54 subcortical parcels from Automated Anatomical Labelling Atlas 3 (AAL3) (Rolls et al., 2020). For each ROI, the timeseries of all voxels were extracted and averaged. Condition-specific FC was estimated for 0-back and 2-back conditions using weighted correlation analysis appropriate for block-design fMRI paradigms (Whitfield-Gabrieli & Nieto-Castanon, 2012). Temporal weights were derived from condition timings (boxcar functions) convolved with canonical hemodynamic response function to account for hemodynamic delays and minimise interference between adjacent task conditions. ROI-to-ROI FC matrices were defined as Fisher-transformed Pearson correlation coefficients representing connectivity strength during each condition (Fisher, 1915; Friston, 2011). Finally, spurious connections were masked out of the FC matrices. Spurious connections were determined as those that have p-value > 0.05. Two-tailed p-values for testing for spurious connections where defined based on the cumulative normal distribution of the Z-statistics.

### Network control theory

Network control theory is a mathematical framework used to examine how a connected system can be steered toward specific target states via external inputs (Kalman, 1963). In the context of neuroscience, brain regions are defined as nodes, and their connections represent the edges of the network. While the approach was initially applied to structural connectivity to investigate how the anatomical architecture constraints brain state transitions (Gu et al., 2015), it has since been extended to FC (Deng et al., 2022; Li et al., 2023). In this functional framework, the FC matrix is used to define the system’s coupling matrix, which governs the assumed linear dynamics of neural activity. This model allows for the derivation of the controllability Gramian, a matrix used to calculate diverse metrics (such as average controllability, modal controllability and activation energy), that quantify a region’s capacity to steer the network into the diverse functional brain states required for cognitive performance (Tang & Bassett, 2018).

#### FC matrix to system matrix transformation

To derive controllability metrics, the dynamics of brain activity are modelled as a simplified, noise-free, linear, continuous-time and time-invariant dynamical system. In this model, the functional brain network is used to define the coupling between regions (the system matrix), allowing us to estimate how activity patterns (or brain states) evolve and transition given the network’s current organisation. That is,

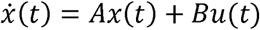

where x (*t*) is the brain state at time *t* (; A is the time-invariant system matrix reflecting interactions between nodes; B is the identity matrix used to specify the nodes used as control inputs; u (*t*) is the input control energy applied at time *t* (. The system matrix A was derived from the FC using a modified Laplacian normalisation (Tang & Bassett, 2018):

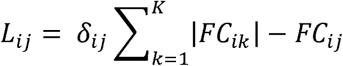

where *FC*_ij_ is the functional connectivity between regions i and j, K is the total number of nodes, and δ_ij_ is the Kronecker delta. The system matrix was finally defined as:

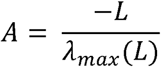

where λ_max_ (L) is the maximum absolute eigenvalue of L, ensuring that the system remains stable (Datta, 2004; Tang & Bassett, 2018).

#### Controllability measures

Average controllability, modal controllability and activation energy were calculated for every node to characterise the controllability of the system.

Average controllability measures a node’s capacity to drive the system to a wide range of easily reachable states with minimal input energy. A brain region with high average controllability acts like a functional hub capable of propagating activity across many regions to drive the network into numerous routine configurations requiring minimal control energy (Gu et al., 2015). Mathematically it is calculated as the trace of the controllability Gramian matrix w_i_ representing the systems reachability when node *i* acts as the external driver:

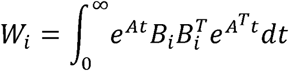

where e^At^ and e^ATt^ are matrix exponentials that describe the evolution and decay of activity patterns across the network’s functional topology over time. The matrix B defines the control set. For regional controllability specifically, Bi is a column vector with a value of one at the i-th position and zero elsewhere, ensuring that node i is the exclusive control input.

Modal controllability measures the capacity of a node to drive the system into difficult-to-reach and high-energy states such as those that require complex, coordinated patterns of activity distinct from baseline dynamics. A brain region with high modal controllability is uniquely positioned to steer brain dynamics towards specialised or non-habitual states, and is thus important for the flexible transition between demanding cognitive states (Gu et al., 2015). It was calculated as:

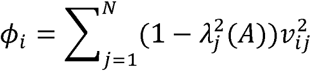

where λ_j_ (A) represents the *j*-th eigenvalue of the system matrix A, which characterises the decay rate of the j-th natural mode of the network; and v= [v_ij_] denotes the eigenvector matrix of A, where the elements v_ij_ quantifies the leverage or “weight” that node i exerts over the j-th mode.

Activation energy quantifies the minimum control energy required to maintain a specific region in an active state, given the surrounding network dynamics. A low activation energy indicates that the region’s activation is naturally supported by the network topology, requiring little external input to remain active. Conversely, high activation energy suggests that the network architecture resists that region’s activity, making its maintenance metabolically expensive and potentially unstable (Gu et al., 2017). It was calculated as:

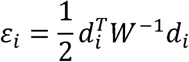

where *d_i_* is a unit vector representing the selective activation of node *i* with value of one at the node and value of zero at all other nodes.

### Between-group comparisons

Between group comparisons were performed to determine whether the NF1 group exhibited differences in performance, and to characterise FC and network controllability in NF1 during the working memory task.

These were performed using general linear models (GLM), which were used to obtain t-statistics for the group effect, while controlling for the confounding effects of age and sex. To determine statistical significance, 10,000 permutation tests were performed to generate a null distribution of t-values. For each permutation, the group labels were randomly shuffled, preserving group sizes, and the GLMs were repeated. Multiple comparisons were corrected using the false discovery rate correction (FDR) method.

These GLMs were specifically fit to behavioural performance measures (accuracy, D-prime, response times and IES) for the working memory task, and region-wise FC values and controllability metrics (average controllability, modal controllability and activation energy). Behavioural performance and FC were also compared for the effect of working memory load (2-back > 0-back) (for FC results see Supplementary Material 2, and Supplementary Tables 6 and 7), while network controllability metrics were compared exclusively for the 2-back condition.

Additionally, between group comparisons were also performed using a network-wise approach to FC and controllability to aid the interpretation of results relative to well recognized functional networks. We repeated GLM modelling for the 17 functional networks individually (Schaefer et al., 2018; Yeo et al., 2011), including Control (A, B, C), Default (A, B, C), Dostal Attention (A, B), Limbic (A, B), Ventral Attention (A, B), Somatomotor (A, B), Temporal Parietal, Central Visual, Peripheral Visual, plus an anatomically defined Subcortical network obtained from AAL3 (Rolls et al., 2020). For FC, network-to-network connectivity was estimated by averaging connections within each network and between each pair of networks, which yielded an 18-by-18 network matrix. For network controllability measures, each nodal metric was averaged over each functional network.

### Classification

Classification was performed with the goal to determine whether control profiles can serve as neural markers of NF1. Specifically, binary classification of neurotypical controls and NF1 participants was performed using linear support vector machine (SVM) classifiers implemented in MATLAB (The MathWorks, Natick, MA) using the fitcsvm function from the Statistics and Machine Learning Toolbox. The classifiers were trained using age, sex, and network controllability measures as predictors. Controllability measures carried into training were those that showed significant group differences in between-group comparisons. Models were trained using only average controllability, only modal controllability, only region activation, and combination of all three. To ensure robust performance estimates, 1,000 repeats of 5-fold cross-validation were performed with sample stratification to preserve group proportions across folds. For each fold, features were standardised using z-score normalisation applied to the training dataset, with the mean and standard deviation of training sample parameters subsequently applied to the test sample. Linear SVM classifiers were then trained on the training sample and applied to predict group membership in the held-out test sample. Classification performance was quantified using balanced accuracy, precision, sensitivity, specificity and area under the receiver operating characteristic curve (AUC), following established guideline for machine learning performance evaluation (Erickson & Kitamura, 2021). Mean performance metrics across all 5,000 evaluations are reported (1,000 repetitions × 5 folds).

### Prediction

The final goal of this work was to use network controllability measures to identify brain regions whose control properties are altered in NF1 and relate to working memory performance. To achieve this goal, support vector regression (SVR) was implemented using a linear kernel to ensure interpretability and prevent overfitting. This method identifies the linear combination of brain features that best predicts working memory performance while preventing overfitting through regularisation. The regularisation parameter C (controlling model complexity) was set to 0.001 to provide strong regularisation and ensure model generalisation, while epsilon (tube width controlling error tolerance) remained at default value 0.1. Other regularising values of parameter C were also explored (0.1, 0.01 and range between 0.0009 and 0.0001), but 0.001 appeared to balance the model fit to training sample and out of sample predictions.

Models were fitted to predict working memory performance in NF1 participants using participant demographics (age, sex), medication status and controllability measures (average controllability, modal controllability, regional energy and combination of all three) from regions that showing significant group differences. Medication status was coded as a binary variable reflecting the use of stimulant, antidepressant and antipsychotic medications expected to affect cognition, brain function, or fMRI signal during scanning (Dichter et al., 2012; Linke et al., 2017; Spencer et al., 2013). During model training, controllability measures and age were standardised using z-score normalisation, while sex and medication status remained binary. Supplementary Material 4 also presents the effects of including effect of motion estimated as mean frame displacement and comorbid diagnoses in the model.

Working memory performance was estimated during model training using principal component analysis (PCA) across multiple tasks (Litwińczuk et al., 2022), capturing variance across in-scanner verbal N-back (d-prime: 0-back and 2-back), out-of-scanner visuospatial N- back (d-prime: 0-back, 1-back, 2-back, and 3-back), and out-of-scanner Corsi Blocks (memory span). This approach addressed the issue of ceiling effects observed in the verbal 2-back condition with many participants attaining high scores. During model training, each behavioural measure was standardised (z-score) and PCA was performed. The resulting first principal component was used to represent working memory performance.

One thousand repeats of 5-fold cross-validation were performed to estimate model generalisability to unseen samples. During cross-validation the normalisation parameters for brain features and task measures, as well as PCA coefficients, were obtained for the training sample and applied to the test sample, while carefully ensuring that no data leakage occurred (Rosenblatt et al., 2024). For each fold of each repeat, models were fitted to the training sample’s data. Model out-of-sample predictive performance was measured for each repeat using Pearson’s R (accuracy of predicted scores) and R^2^ (explained variance) (Valente et al., 2021). Following cross-validation, models were fitted to the complete sample of NF1 participants to estimate final model fit and obtain interpretable beta coefficients (feature weights) (Tsamardinos et al., 2018).

Permutation testing was performed to estimate the significance of the model’s predictive performance during cross-validation (R^2^), and to estimate the significance of the beta coefficients in the final model. During 10,000 permutations, each working memory variable was shuffled across subjects and models were refitted, generating null models. The predictive performance of the empirical models was compared to that of null models to determine the proportion of permutations where the observed R^2^ exceeded the null R^2^ (one-tailed) (Phipson & Smyth, 2010). For beta coefficients, two-tailed p-values were calculated as twice the proportion of permutations where observed absolute values exceeded the null absolute values.

### Visualisations

All the FC networks presented in this report were visualized with the BrainNet Viewer (http://www.nitrc.org/projects/bnv/) (Xia et al., 2013). Region-specific projections of controllability presented in this report were visualized using MRIcroGL (https://www.nitrc.org/projects/mricrogl/), and Nilearn (0.13.1) and Matplotlib (v3.10.9) in Python (3.10.9).

## Supporting information

Supplementary Results

## Acknowledgements

This research was supported by the NIHR Manchester Biomedical Research Centre (NIHR203308). ML was funded by the Office for Life Sciences and the National Institute for Health and Care Research (NIHR) Mental Health Translational Research Collaboration, hosted by the NIHR Oxford Health Biomedical Research Centre (NIHR203308). The views expressed are those of the author(s) and not necessarily those of the NIHR or the Department of Health and Social Care. The authors also wish to thank the NF1 participants and families that participated in this study. This work was also supported by the Neurofibromatosis Therapeutic Acceleration Program (NTAP) through a Francis Collins Scholarship to SG. NT is supported by Medical Research Council (MRC) (MR/X005267/1). JJ was supported by AMS Springboard (SBF007\100077) and the MRC Programme (UKRI527). JY is supported by US Department of Defence (NF230043)

## Data/code availability

The NF1 participant data have been deposited on the Sage Bionetworks data repository https://www.synapse.org/. Approved researchers can request to obtain the data which are subject to data sharing agreements.

## Ethics statement

Ethics approval for the study was obtained from the North West-Greater Manchester South Research Ethics Committee (reference: 18/NW/0762). Written informed consent was obtained from the parents and older adolescent participants and assent was obtained from the younger participants.

## References

Ashburner, J. (2007). A fast diffeomorphic image registration algorithm. NeuroImage, 38(1), 95–113. 10.1016/j.neuroimage.2007.07.007

Baddeley, A. (2003). Working memory: looking back and looking forward. Nature Reviews Neuroscience, 4(10), 829–839. 10.1038/nrn1201

Baddeley, A. (2012). Working Memory: Theories, Models, and Controversies. Annual Review of Psychology, 63(1), 1–29. 10.1146/annurev-psych-120710-100422

Barbey, A. K., Koenigs, M., & Grafman, J. (2013). Dorsolateral prefrontal contributions to human working memory. Cortex, 49(5), 1195–1205. 10.1016/j.cortex.2012.05.022

Bassett, D. S., Wymbs, N. F., Porter, M. A., Mucha, P. J., Carlson, J. M., & Grafton, S. T. (2011). Dynamic reconfiguration of human brain networks during learning. Proceedings of the National Academy of Sciences, 108(18), 7641–7646. 10.1073/pnas.1018985108

Baudou, E., Nemmi, F., Biotteau, M., Maziero, S., Peran, P., & Chaix, Y. (2020). Can the Cognitive Phenotype in Neurofibromatosis Type 1 (NF1) Be Explained by Neuroimaging? A Review. Frontiers in Neurology, 10. 10.3389/fneur.2019.01373

Behzadi, Y., Restom, K., Liau, J., & Liu, T. T. (2007). A component based noise correction method (CompCor) for BOLD and perfusion based fMRI. NeuroImage, 37(1), 90–101. 10.1016/j.neuroimage.2007.04.042

Beynel, L., Deng, L., Crowell, C. A., Dannhauer, M., Palmer, H., Hilbig, S., Peterchev, A. V., Luber, B., Lisanby, S. H., Cabeza, R., Appelbaum, L. G., & Davis, S. W. (2020). Structural Controllability Predicts Functional Patterns and Brain Stimulation Benefits Associated with Working Memory. The Journal of Neuroscience, 40(35), 6770–6778. 10.1523/jneurosci.0531-20.2020

Bruyer, R., & Brysbaert, M. (2011). Combining Speed and Accuracy in Cognitive Psychology: Is the Inverse Efficiency Score (IES) a Better Dependent Variable than the Mean Reaction Time (RT) and the Percentage Of Errors (PE)? Psychologica Belgica, 51(1). 10.5334/pb-51-1-5

Corbetta, M., & Shulman, G. L. (2002). Control of goal-directed and stimulus-driven attention in the brain. Nature Reviews Neuroscience, 3(3), 201–215. 10.1038/nrn755

Costa, R. M., Federov, N. B., Kogan, J. H., Murphy, G. G., Stern, J., Ohno, M., Kucherlapati, R., Jacks, T., & Silva, A. J. (2002). Mechanism for the learning deficits in a mouse model of neurofibromatosis type 1. Nature, 415(6871), 526–530. 10.1038/nature711

Datta, B. N. (2004). Numerical Methods for Linear Control Systems. 10.1016/b978-0-12-203590-6.X5000-9

Deng, S., Li, J., Thomas Yeo, B. T., & Gu, S. (2022). Control theory illustrates the energy efficiency in the dynamic reconfiguration of functional connectivity. Communications Biology, 5(1). 10.1038/s42003-022-03196-0

Dichter, G. S., Sikich, L., Song, A., Voyvodic, J., & Bodfish, J. W. (2012). Functional Neuroimaging of Treatment Effects in Psychiatry: Methodological Challenges and Recommendations. International Journal of Neuroscience, 122(9), 483–493. 10.3109/00207454.2012.678446

Dipasquale, O., Martins, D., Sethi, A., Veronese, M., Hesse, S., Rullmann, M., Sabri, O., Turkheimer, F., Harrison, N. A., Mehta, M. A., & Cercignani, M. (2020). Unravelling the effects of methylphenidate on the dopaminergic and noradrenergic functional circuits. Neuropsychopharmacology, 45(9), 1482–1489. 10.1038/s41386-020-0724-x

Erickson, B. J., & Kitamura, F. (2021). Magician’s Corner: 9. Performance Metrics for Machine Learning Models. Radiology: Artificial Intelligence, 3(3). 10.1148/ryai.2021200126

Evans, D. G., Howard, E., Giblin, C., Clancy, T., Spencer, H., Huson, S. M., & Lalloo, F. (2010). Birth incidence and prevalence of tumor-prone syndromes: Estimates from a UK family genetic register service. American Journal of Medical Genetics Part A, *152A*(2), 327-332. 10.1002/ajmg.a.33139

Fisher, R. A. (1915). Frequency Distribution of the Values of the Correlation Coefficient in Samples from an Indefinitely Large Population. Biometrika, 10(4). 10.2307/2331838

Friston, K. J. (2011). Functional and Effective Connectivity: A Review. Brain Connectivity, 1(1), 13–36. 10.1089/brain.2011.0008

Garg, S., Williams, S., Jung, J., Pobric, G., Nandi, T., Lim, B., Vassallo, G., Green, J., Evans, D. G., Stagg, C. J., Parkes, L. M., & Stivaros, S. (2022). Non-invasive brain stimulation modulates GABAergic activity in neurofibromatosis 1. Scientific Reports, 12(1). 10.1038/s41598-022-21907-9

Gu, S., Betzel, R. F., Mattar, M. G., Cieslak, M., Delio, P. R., Grafton, S. T., Pasqualetti, F., & Bassett, D. S. (2017). Optimal trajectories of brain state transitions. NeuroImage, 148, 305–317. 10.1016/j.neuroimage.2017.01.003

Gu, S., Pasqualetti, F., Cieslak, M., Telesford, Q. K., Yu, A. B., Kahn, A. E., Medaglia, J. D., Vettel, J. M., Miller, M. B., Grafton, S. T., & Bassett, D. S. (2015). Controllability of structural brain networks. Nature Communications, 6(1). 10.1038/ncomms9414

Gutmann, D. H., Ferner, R. E., Listernick, R. H., Korf, B. R., Wolters, P. L., & Johnson, K. J. (2017). Neurofibromatosis type 1. Nature Reviews Disease Primers, 3(1). 10.1038/nrdp.2017.4

Halai, A. D., Welbourne, S. R., Embleton, K., & Parkes, L. M. (2014). A comparison of dual gradient-echo and spin-echo fMRI of the inferior temporal lobe. Human Brain Mapping, 35(8), 4118–4128. 10.1002/hbm.22463

Huntenburg, J. M., Bazin, P.-L., & Margulies, D. S. (2018). Large-Scale Gradients in Human Cortical Organization. Trends in Cognitive Sciences, 22(1), 21–31. 10.1016/j.tics.2017.11.002

Ibrahim, A. F. A., Montojo, C. A., Haut, K. M., Karlsgodt, K. H., Hansen, L., Congdon, E., Rosser, T., Bilder, R. M., Silva, A. J., & Bearden, C. E. (2017). Spatial working memory in neurofibromatosis 1: Altered neural activity and functional connectivity. NeuroImage: Clinical, 15, 801–811. 10.1016/j.nicl.2017.06.032

Jaeggi, S. M., Studer-Luethi, B., Buschkuehl, M., Su, Y.-F., Jonides, J., & Perrig, W. J. (2010). The relationship between n-back performance and matrix reasoning — implications for training and transfer. Intelligence, 38(6), 625–635. 10.1016/j.intell.2010.09.001

Kalman, R. (1963). Controllability of linear dynamical systems. Contributions to differential equations, 189-213.

Kirchner, W. K. (1958). Age differences in short-term retention of rapidly changing information. Journal of Experimental Psychology, 55(4), 352–358. 10.1037/h0043688

Lehtonen, A., Howie, E., Trump, D., & Huson, S. M. (2012). Behaviour in children with neurofibromatosis type 1: cognition, executive function, attention, emotion, and social competence. Developmental Medicine & Child Neurology, 55(2), 111–125. 10.1111/j.1469-8749.2012.04399.x

Li, Q., Yao, L., You, W., Liu, J., Deng, S., Li, B., Luo, L., Zhao, Y., Wang, Y., Wang, Y., Zhang, Q., Long, F., Sweeney, J. A., Gu, S., Li, F., & Gong, Q. (2023). Controllability of Functional Brain Networks and Its Clinical Significance in First-Episode Schizophrenia. Schizophrenia Bulletin, 49(3), 659–668. 10.1093/schbul/sbac177

Linke, A. C., Olson, L., Gao, Y., Fishman, I., & Müller, R.-A. (2017). Psychotropic Medication Use in Autism Spectrum Disorders May Affect Functional Brain Connectivity. Biological Psychiatry: Cognitive Neuroscience and Neuroimaging, 2(6), 518–527. 10.1016/j.bpsc.2017.06.008

Litwińczuk, M. C., Garg, S., Lea-Carnall, C., & Trujillo-Barreto, N. J. (2026). Disrupted Frontoparietal Dynamics in Neurofibromatosis Type 1: Reduced Sensitivity and Atypical Modulation During Working Memory. Human Brain Mapping, 47(2). 10.1002/hbm.70464

Litwińczuk, M. C., Garg, S., Williams, S. R., Green, J., Lea-Carnall, C., & Trujillo-Barreto, N. J. (2025). Non-invasive brain stimulation reorganises effective connectivity during a working memory task in individuals with Neurofibromatosis Type 1. NeuroImage: Reports, 5(2). 10.1016/j.ynirp.2025.100258

Litwińczuk, M. C., Muhlert, N., Cloutman, L., Trujillo-Barreto, N., & Woollams, A. (2022). Combination of structural and functional connectivity explains unique variation in specific domains of cognitive function. NeuroImage, 262. 10.1016/j.neuroimage.2022.119531

Macmillan, N. A., & Creelman, C. D. (2004). Detection Theory. 10.4324/9781410611147

Margulies, D. S., Ghosh, S. S., Goulas, A., Falkiewicz, M., Huntenburg, J. M., Langs, G., Bezgin, G., Eickhoff, S. B., Castellanos, F. X., Petrides, M., Jefferies, E., & Smallwood, J. (2016). Situating the default-mode network along a principal gradient of macroscale cortical organization. Proceedings of the National Academy of Sciences, 113(44), 12574–12579. 10.1073/pnas.1608282113

Mennigen, E., Schuette, P., Vajdi, A., Pacheco, L., Rosser, T., & Bearden, C. E. (2019). Reduced higher dimensional temporal dynamism in neurofibromatosis type 1. NeuroImage: Clinical, 22. 10.1016/j.nicl.2019.101692

Mesulam, M. (1998). From sensation to cognition. Brain, 121(6), 1013–1052. 10.1093/brain/121.6.1013

Mesulam, M. (2008). Representation, inference, and transcendent encoding in neurocognitive networks of the human brain. Annals of Neurology, 64(4), 367–378. 10.1002/ana.21534

Nieto-Castanon, A. (2020). FMRI denoising pipeline. In Handbook of functional connectivity Magnetic Resonance Imaging methods in CONN (pp. 17-25). 10.56441/hilbertpress.2207.6600

Owen, A. M., McMillan, K. M., Laird, A. R., & Bullmore, E. (2005). N-back working memory paradigm: A meta-analysis of normative functional neuroimaging studies. Human Brain Mapping, 25(1), 46–59. 10.1002/hbm.20131

Phipson, B., & Smyth, G. K. (2010). Permutation P-values Should Never Be Zero: Calculating Exact P-values When Permutations Are Randomly Drawn. Statistical Applications in Genetics and Molecular Biology, 9(1). 10.2202/1544-6115.1585

Plasschaert, E., Van Eylen, L., Descheemaeker, M. J., Noens, I., Legius, E., & Steyaert, J. (2016). Executive functioning deficits in children with neurofibromatosis type 1: The influence of intellectual and social functioning. American Journal of Medical Genetics Part B: Neuropsychiatric Genetics, 171(3), 348–362. 10.1002/ajmg.b.32414

Pobric, G., Taylor, J. R., Ramalingam, H. M., Pye, E., Robinson, L., Vassallo, G., Jung, J., Bhandary, M., Szumanska-Ryt, K., Theodosiou, L., Evans, D. G., Eelloo, J., Burkitt-Wright, E., Hulleman, J., Green, J., & Garg, S. (2021). Cognitive and Electrophysiological Correlates of Working Memory Impairments in Neurofibromatosis Type 1. Journal of Autism and Developmental Disorders, 52(4), 1478–1494. 10.1007/s10803-021-05043-3

Power, J. D., Barnes, K. A., Snyder, A. Z., Schlaggar, B. L., & Petersen, S. E. (2012). Spurious but systematic correlations in functional connectivity MRI networks arise from subject motion. NeuroImage, 59(3), 2142–2154. 10.1016/j.neuroimage.2011.10.018

Raichle, M. E., MacLeod, A. M., Snyder, A. Z., Powers, W. J., Gusnard, D. A., & Shulman, G. L. (2001). A default mode of brain function. Proceedings of the National Academy of Sciences, 98(2), 676–682. 10.1073/pnas.98.2.676

Rolls, E. T., Huang, C.-C., Lin, C.-P., Feng, J., & Joliot, M. (2020). Automated anatomical labelling atlas 3. NeuroImage, 206. 10.1016/j.neuroimage.2019.116189

Rosenblatt, M., Tejavibulya, L., Jiang, R., Noble, S., & Scheinost, D. (2024). Data leakage inflates prediction performance in connectome-based machine learning models. Nature Communications, 15(1). 10.1038/s41467-024-46150-w

Saberi, M., Rieck, J. R., Golafshan, S., Grady, C. L., Misic, B., Dunkley, B. T., & Khatibi, A. (2024). The brain selectively allocates energy to functional brain networks under cognitive control. Scientific Reports, 14(1). 10.1038/s41598-024-83696-7

Sawyer, C., Green, J., Lim, B., Pobric, G., Jung, J., Vassallo, G., Evans, D. G., Stagg, C. J., Parkes, L. M., Stivaros, S., Muhlert, N., & Garg, S. (2022). Neuroanatomical correlates of working memory performance in Neurofibromatosis 1. Cerebral Cortex Communications, 3(2). 10.1093/texcom/tgac021

Schaefer, A., Kong, R., Gordon, E. M., Laumann, T. O., Zuo, X.-N., Holmes, A. J., Eickhoff, S. B., & Yeo, B. T. T. (2018). Local-Global Parcellation of the Human Cerebral Cortex from Intrinsic Functional Connectivity MRI. Cerebral Cortex, 28(9), 3095–3114. 10.1093/cercor/bhx179

Shilyansky, C., Karlsgodt, K. H., Cummings, D. M., Sidiropoulou, K., Hardt, M., James, A. S., Ehninger, D., Bearden, C. E., Poirazi, P., Jentsch, J. D., Cannon, T. D., Levine, M. S., & Silva, A. J. (2010). Neurofibromin regulates corticostriatal inhibitory networks during working memory performance. Proceedings of the National Academy of Sciences, 107(29), 13141–13146. 10.1073/pnas.1004829107

Spencer, T. J., Brown, A., Seidman, L. J., Valera, E. M., Makris, N., Lomedico, A., Faraone, S. V., & Biederman, J. (2013). Effect of Psychostimulants on Brain Structure and Function in ADHD. The Journal of Clinical Psychiatry, 74(09), 902–917. 10.4088/JCP.12r08287

Swanson, H. L., & Alloway, T. P. (2012). Working memory, learning, and academic achievement. In APA educational psychology handbook, Vol 1: Theories, constructs, and critical issues. (pp. 327-366). 10.1037/13273-012

Tang, E., & Bassett, D. S. (2018). Colloquium: Control of dynamics in brain networks. Reviews of Modern Physics, 90(3). 10.1103/RevModPhys.90.031003

Tsamardinos, I., Greasidou, E., & Borboudakis, G. (2018). Bootstrapping the out-of-sample predictions for efficient and accurate cross-validation. Machine Learning, 107(12), 1895–1922. 10.1007/s10994-018-5714-4

Tu, C., Rocha, R. P., Corbetta, M., Zampieri, S., Zorzi, M., & Suweis, S. (2018). Warnings and caveats in brain controllability. NeuroImage, 176, 83–91. 10.1016/j.neuroimage.2018.04.010

Valente, G., Castellanos, A. L., Hausfeld, L., De Martino, F., & Formisano, E. (2021). Cross-validation and permutations in MVPA: Validity of permutation strategies and power of cross-validation schemes. NeuroImage, 238. 10.1016/j.neuroimage.2021.118145

Violante, I. R., Patricio, M., Bernardino, I., Rebola, J., Abrunhosa, A. J., Ferreira, N., & Castelo-Branco, M. (2016). GABA deficiency in NF1. Neurology, 87(9), 897–904. 10.1212/wnl.0000000000003044

Violante, I. R., Ribeiro, M. J., Cunha, G., Bernardino, I., Duarte, J. V., Ramos, F., Saraiva, J., Silva, E., & Castelo-Branco, M. (2012). Abnormal Brain Activation in Neurofibromatosis Type 1: A Link between Visual Processing and the Default Mode Network. PLoS ONE, 7(6). 10.1371/journal.pone.0038785

Wang, F., Liu, Z., Wang, J., Li, X., Pan, Y., Yang, J., Cheng, P., Sun, F., Tan, W., Huang, D., Zhang, J., Liu, X., Zhong, M., Wu, G., Yang, J., & Palaniyappan, L. (2025). Aberrant controllability of functional connectome during working memory tasks in patients with schizophrenia and unaffected siblings. The British Journal of Psychiatry, 227(1), 463–472. 10.1192/bjp.2024.225

Whitfield-Gabrieli, S., & Nieto-Castanon, A. (2012). Conn: A Functional Connectivity Toolbox for Correlated and Anticorrelated Brain Networks. Brain Connectivity, 2(3), 125–141. 10.1089/brain.2012.0073

Wild, A., Tziraki, M., Lea-Carnall, C., & Garg, S. (2026). A Systematic Review of Functional Brain Imaging Studies in Neurofibromatosis 1. Neuropsychology Review. 10.1007/s11065-026-09695-9

Xia, M., Wang, J., & He, Y. (2013). BrainNet Viewer: A Network Visualization Tool for Human Brain Connectomics. PLoS ONE, 8(7). 10.1371/journal.pone.0068910

Yamashita, M., Yoshihara, Y., Hashimoto, R., Yahata, N., Ichikawa, N., Sakai, Y., Yamada, T., Matsukawa, N., Okada, G., Tanaka, S. C., Kasai, K., Kato, N., Okamoto, Y., Seymour, B., Takahashi, H., Kawato, M., & Imamizu, H. (2018). A prediction model of working memory across health and psychiatric disease using whole-brain functional connectivity. eLife, 7. 10.7554/eLife.38844

Yang, J., Liu, Z., Wang, F., Tan, W., Huang, D., Ouyang, X., Tao, H., Wu, G., Pan, Y., Yang, J., & Palaniyappan, L. (2025). Task-Related Controllability of Functional Connectome During a Working Memory Task in Schizophrenia, Bipolar Disorder, and Major Depressive Disorder. Research, 8. 10.34133/research.0792

Yang, Y., Woollams, A., Lipp, I., Zhuo, Z., Litwińczuk, M. C., Tomassini, V., Liu, Y., Trujillo-Barreto, N. J., & Muhlert, N. (2025). Thalamic Network Controllability Predicts Cognitive Impairment in Multiple Sclerosis. Human Brain Mapping, 46(10). 10.1002/hbm.70284

Yeo, T. B. T., Krienen, F. M., Sepulcre, J., Sabuncu, M. R., Lashkari, D., Hollinshead, M., Roffman, J. L., Smoller, J. W., Zöllei, L., Polimeni, J. R., Fischl, B., Liu, H., & Buckner, R. L. (2011). The organization of the human cerebral cortex estimated by intrinsic functional connectivity. Journal of Neurophysiology, 106(3), 1125–1165. 10.1152/jn.00338.2011

