## Supplementary Results for "Working memory impairments in Neurofibromatosis Type 1 are explained by disrupted functional connectivity and network controllability"

### 1. Out-of-scanner behavioural group differences

Supplementary Table 1 presents the descriptive statistics of group performance during Visuo-spatial N-back task and Corsi Blocks task. During the visuo-spatial N-back task, the NF1 group showed no significant difference from the control group in 0-back condition (t(84)=-0.04, p=0.9647, β =-0.0381), but showed significantly lower accuracy during 1-back (t(84)=-2.94, p=0.0042, β =-3.7235), 2-back (t(84)=-5.05, p=0.0000, β =-10.7107), and 3-back conditions (t(84)=-3.04, p=0.0032, β =-5.8761). Similarly, for d-prime there was no significant difference during 0-back condition (t(84)=-1.62, p=0.1090, β =-0.2015), but the NF1 group had significantly lower d-prime during 1-back (t(84)=-3.94, p=0.0002, β =-0.6027), 2-back (t(84)=-5.44, p=0.0000, β =-1.1033), and 3-back (t(84)=-2.93, p=0.0044, β =-0.4619) conditions. No significant group differences in response times were observed for the 0-back (t(84) = 1.74, p = .085, β = 28.60), 1-back (t(84) = 0.45, p = .652, β = 13.84), or 2-back conditions (t(84) = -1.52, p = .131, β = -68.65). During 3-back condition, the NF1 group showed significantly lower response times than controls (t(84) = -3.35, p = .001, β = -201.50). There were not significant group differences in IES during 0-back (t(84) = 1.69, p = .096, β = 29.05), 1-back (t(84) = 1.13, p = .262, β = 39.50), or 2-back conditions (t(84) = 0.20, p = .841, β = 13.21). During 3-back condition, the NF1 group showed significantly lower IES than controls (t(84) = -2.52, p = .014, β = -213.35). During the Corsi Blocks task, the NF1 group showed significantly lower baseline memory span than controls (t(82) = -6.28, p = 0.000, β = -1.25).

| Supplementary Table 1 The descriptive statistics of group performance on baseline tasks. | | | | | |
| --- | --- | --- | --- | --- | --- |
| **Measure** | **Task condition** | **Control group** | | **NF1 group** | |
|  |  | Mean | SD | Mean | SD |
| Accuracy (%) | 0-back | 97.18 | 3.012 | 97.192 | 4.521 |
|  | 1-back | 95.194 | 3.117 | 91.469 | 7.524 |
|  | 2-back | 90.32 | 7.754 | 79.779 | 10.998 |
|  | 3-back | 76.49 | 8.888 | 70.637 | 8.755 |
| Response time (ms) | 0-back | 372.812 | 63.545 | 399.559 | 88.15 |
|  | 1-back | 480.298 | 137.278 | 494.195 | 146.944 |
|  | 2-back | 651.433 | 222.053 | 581.069 | 198.276 |
|  | 3-back | 803.702 | 338.459 | 601.949 | 231.957 |
| D-Prime | 0-back | 4.005 | 0.394 | 3.808 | 0.669 |
|  | 1-back | 3.584 | 0.484 | 2.982 | 0.849 |
|  | 2-back | 2.922 | 0.95 | 1.839 | 0.928 |
|  | 3-back | 1.501 | 0.776 | 1.037 | 0.675 |
| Inverse efficiency score | 0-back | 384.905 | 71.495 | 411.887 | 91.146 |
|  | 1-back | 505.179 | 146.321 | 545.258 | 178.41 |
|  | 2-back | 736.94 | 291.111 | 748 | 310.681 |
|  | 3-back | 1071.233 | 476.029 | 858.259 | 329.697 |
| Memory span | Corsi Blocks | 5.743 | 0.852 | 4.5 | 0.943 |

### 2. Functional Connectivity

#### 2.1. 2-back>0-back contrast

Following false discovery rate correction for multiple comparisons, increased connectivity in NF1 sample between left lateral prefrontal cortex (control A) and right somatomotor cortex (somatomotor A) was shared between 2-back contrast and 2-back>0-back contrast. The two contrasts also shared increased connectivity in NF1 between different region of dorsal attention and somatomotor networks, with the 2-back>0-back contrast showing increased connectivity between left somatomotor cortex and left dorsal prefrontal cortex (default A), and left ventral prefrontal cortex (default B) and left temporal parietal cortex. Uniquely, 2-back>0-back showed increased connectivity between left somatomotor cortex (somatomotor A) and left dorsal prefrontal cortex (default A), and between left ventral prefrontal cortex (default B) and left temporal parietal cortex.

#### 2.2. Using 2-back FC as predictors

Predictive modelling was repeated using FDR-corrected 2-back functional connectivity (n = 37) as predictors of composite working memory component. The model achieved a cross-validation R² = 0.015, which showed a trend toward significance (null R² = -0.053 ± 0.058, *p* = 0.075). The final model fitted to the whole NF1 sample achieved R² = 0.030.

Combination of FC and all controllability measures showed poorer predictive accuracy than use of controllability measures alone (cross-validation R² = 0.006).

### 3. Network Control Theory

#### 3.1. PCA identifies a general working memory component

Because the behavioural prediction hypothesis concerned individual differences in general working memory ability rather than performance on a single task, we first derived a composite measure of working memory using PCA. The first component explained 45.9% of variance in behavioural performance of all 51 NF1 participants. During cross-validation, it explained 46.0% ± 3.2% of variance across 5000 models (1,000 repetitions × 5-fold cross-validation). The coefficients of the first component (Supplementary Table 2) were largest for the 1-back and 2-back working memory load regardless of verbal or visuospatial modality, therefore they appeared to describe a general working memory component. This component was carried to further predictive modelling.

| Supplementary Table 2. Coefficients of the first component generated by the PCA analysis of working memory tasks. Cross-validation values represent the mean ± standard deviations in PCA coefficients across the 5000 models (1,000 repetitions × 5-fold cross-validation). | | |
| --- | --- | --- |
| Task | Final model coefficients | Cross-validation |
| Verbal 0-back | 0.353 | 0.349 ± 0.034 |
| Verbal 2-back | 0.418 | 0.417 ± 0.014 |
| Visuospatial 0-back | 0.331 | 0.328 ± 0.037 |
| Visuospatial 1-back | 0.407 | 0.406 ± 0.019 |
| Visuospatial 2-back | 0.455 | 0.455 ± 0.018 |
| Visuospatial 3-back | 0.360 | 0.359 ± 0.023 |
| Corsi Blocks | 0.298 | 0.300 ± 0.045 |

#### 3.2. Predictive probability value of empirical and null models of network controllability

We examined whether controllability measures showing significant group differences between NF1 participants and neurotypical controls could predict individual differences in working memory performance among NF1 participants. Out-of-sample predictive performance was evaluated with 1,000 repetitions of cross-validation, using controllability measures as predictors of the first PCA component. The average controllability model explained a statistically significant portion of variance in the general working memory component (R^2^ = 0.022 ± 0.018), compared with a null distribution (null R² = −0.053 ± 0.058, p = 0.049). The modal controllability model (R² = 0.017 ± 0.018, null R² = −0.053 ± 0.059, p = 0.064), activation energy model (R² = 0.018 ± 0.018, null R² = −0.053 ± 0.058, p = 0.058), and combined model (R² = 0.014 ± 0.018, null R² = −0.061 ± 0.066, p = 0.094) did not reach statistical significance.

In the models fitted to the complete sample of 51 NF1 participants, activation energy showed the greatest predictive performance among the individual metrics (R^2^ = 0.034), while the combined model showed the highest overall model fit (R^2^ = 0.072).

#### 3.3. Predictive models - beta weights

| Supplementary Table 3. All beta weights from the final predictive models of behaviour (n = 51) that pass the statistical *p*-value threshold of 0.05. Parcel indices, network labels, and full labels originate directly from the Schafer et al (2018) atlas. | | | | |
| --- | --- | --- | --- | --- |
| Parcel index | Network | Full label | β | p-value |
| ***Average controllability model*** | | | | |
| 264 | ContB | RH_ContB_PFClv_3 | -0.0168 | <0.001 |
| 104 | ContB | LH_ContB_PFClv_1 | -0.0119 | 0.004 |
| 108 | ContC | LH_ContC_pCun_2 | -0.0112 | 0.004 |
| 54 | DorsAttnA | LH_DorsAttnA_SPL_3 | -0.0105 | 0.006 |
| 133 | DefaultB | LH_DefaultB_PFCl_1 | -0.0103 | 0.008 |
| 291 | DefaultC | RH_DefaultC_Rsp_1 | +0.0095 | 0.019 |
| 149 | TempPar | LH_TempPar_4 | +0.0090 | 0.016 |
| 260 | ContB | RH_ContB_PFCld_2 | -0.0086 | 0.021 |
| 65 | SalVentAttnA | LH_SalVentAttnA_ParOper_2 | -0.0074 | 0.041 |
| 228 | SalVentAttnB | RH_SalVentAttnB_PFCl_1 | -0.0074 | 0.046 |
| 245 | ContA | RH_ContA_Temp_1 | -0.0068 | 0.059 |
| 255 | ContB | RH_ContB_Temp_1 | -0.0061 | 0.082 |
| 150 | TempPar | LH_TempPar_5 | +0.0057 | 0.105 |
| 186 | SomMotB | RH_SomMotB_Aud_1 | -0.0055 | 0.120 |
| 53 | DorsAttnA | LH_DorsAttnA_SPL_2 | -0.0055 | <0.001 |
| 148 | TempPar | LH_TempPar_3 | +0.0053 | <0.001 |
| 320 | ThalVPL | subcortical_ThalVPL_130 | -0.0051 | <0.001 |
| 63 | DorsAttnB | LH_DorsAttnB_FEF_2 | -0.0046 | <0.001 |
| 159 | VisCent | RH_VisCent_ExStr_8 | -0.0042 | <0.001 |
| 262 | ContB | RH_ContB_PFClv_1 | +0.0030 | <0.001 |
| 119 | DefaultA | LH_DefaultA_PFCm_1 | +0.0025 | <0.001 |
| 268 | ContC | RH_ContC_pCun_2 | +0.0024 | <0.001 |
| 235 | LimbicB | RH_LimbicB_OFC_2 | +0.0022 | <0.001 |
| 258 | ContB | RH_ContB_IPL_3 | -0.0022 | <0.001 |
| 123 | DefaultB | LH_DefaultB_Temp_2 | -0.0020 | <0.001 |
| 92 | ContA | LH_ContA_IPS_3 | +0.0019 | <0.001 |
| 192 | SomMotB | RH_SomMotB_S2_5 | +0.0018 | <0.001 |
| 155 | VisCent | RH_VisCent_ExStr_5 | +0.0015 | <0.001 |
| 80 | LimbicB | LH_LimbicB_OFC_1 | +0.0005 | <0.001 |
| 112 | DefaultA | LH_DefaultA_PFCd_1 | -0.0004 | <0.001 |
| ***Modal controllability model*** | | | | |
| 104 | ContB | LH_ContB_PFClv_1 | +0.0117 | 0.003 |
| 54 | DorsAttnA | LH_DorsAttnA_SPL_3 | +0.0106 | 0.006 |
| 108 | ContC | LH_ContC_pCun_2 | +0.0103 | 0.007 |
| 101 | ContB | LH_ContB_Temp_1 | +0.0098 | 0.015 |
| 260 | ContB | RH_ContB_PFCld_2 | +0.0094 | 0.017 |
| 149 | TempPar | LH_TempPar_4 | -0.0089 | 0.019 |
| 128 | DefaultB | LH_DefaultB_PFCd_1 | +0.0086 | 0.026 |
| 291 | DefaultC | RH_DefaultC_Rsp_1 | -0.0085 | 0.029 |
| 148 | TempPar | LH_TempPar_3 | -0.0077 | 0.044 |
| 245 | ContA | RH_ContA_Temp_1 | +0.0073 | 0.056 |
| 228 | SalVentAttnB | RH_SalVentAttnB_PFCl_1 | +0.0068 | 0.063 |
| 150 | TempPar | LH_TempPar_5 | -0.0067 | 0.069 |
| 255 | ContB | RH_ContB_Temp_1 | +0.0066 | 0.073 |
| 65 | SalVentAttnA | LH_SalVentAttnA_ParOper_2 | +0.0061 | 0.083 |
| 186 | SomMotB | RH_SomMotB_Aud_1 | +0.0050 | <0.001 |
| 320 | ThalVPL | subcortical_ThalVPL_130 | +0.0048 | <0.001 |
| 53 | DorsAttnA | LH_DorsAttnA_SPL_2 | +0.0047 | <0.001 |
| 141 | DefaultC | LH_DefaultC_Rsp_1 | +0.0046 | <0.001 |
| 49 | DorsAttnA | LH_DorsAttnA_TempOcc_2 | +0.0045 | <0.001 |
| 325 | ThalMDm | subcortical_ThalMDm_135 | -0.0039 | <0.001 |
| 123 | DefaultB | LH_DefaultB_Temp_2 | +0.0039 | <0.001 |
| 159 | VisCent | RH_VisCent_ExStr_8 | +0.0033 | <0.001 |
| 268 | ContC | RH_ContC_pCun_2 | -0.0033 | <0.001 |
| 92 | ContA | LH_ContA_IPS_3 | -0.0029 | <0.001 |
| 262 | ContB | RH_ContB_PFClv_1 | -0.0029 | <0.001 |
| 192 | SomMotB | RH_SomMotB_S2_5 | -0.0026 | <0.001 |
| 155 | VisCent | RH_VisCent_ExStr_5 | -0.0015 | <0.001 |
| 112 | DefaultA | LH_DefaultA_PFCd_1 | -0.0014 | <0.001 |
| 119 | DefaultA | LH_DefaultA_PFCm_1 | -0.0009 | <0.001 |
| 235 | LimbicB | RH_LimbicB_OFC_2 | -0.0004 | <0.001 |
| 80 | LimbicB | LH_LimbicB_OFC_1 | +0.0001 | <0.001 |
| ***Activation energy model*** | | | | |
| 104 | ContB | LH_ContB_PFClv_1 | +0.0124 | 0.002 |
| 89 | ContA | LH_ContA_Temp_1 | +0.0109 | 0.006 |
| 101 | ContB | LH_ContB_Temp_1 | +0.0107 | 0.008 |
| 142 | DefaultC | LH_DefaultC_Rsp_2 | -0.0102 | 0.012 |
| 54 | DorsAttnA | LH_DorsAttnA_SPL_3 | +0.0100 | 0.009 |
| 255 | ContB | RH_ContB_Temp_1 | +0.0098 | 0.013 |
| 128 | DefaultB | LH_DefaultB_PFCd_1 | +0.0084 | 0.026 |
| 148 | TempPar | LH_TempPar_3 | -0.0081 | 0.035 |
| 243 | LimbicA | RH_LimbicA_TempPole_5 | +0.0080 | 0.034 |
| 245 | ContA | RH_ContA_Temp_1 | +0.0074 | 0.044 |
| 291 | DefaultC | RH_DefaultC_Rsp_1 | -0.0070 | 0.061 |
| 268 | ContC | RH_ContC_pCun_2 | -0.0069 | 0.064 |
| 333 | ThalPuA | subcortical_ThalPuA_143 | -0.0066 | 0.067 |
| 108 | ContC | LH_ContC_pCun_2 | +0.0062 | 0.087 |
| 149 | TempPar | LH_TempPar_4 | -0.0059 | <0.001 |
| 90 | ContA | LH_ContA_IPS_1 | +0.0057 | <0.001 |
| 123 | DefaultB | LH_DefaultB_Temp_2 | +0.0055 | <0.001 |
| 48 | DorsAttnA | LH_DorsAttnA_TempOcc_1 | +0.0054 | <0.001 |
| 186 | SomMotB | RH_SomMotB_Aud_1 | +0.0050 | <0.001 |
| 150 | TempPar | LH_TempPar_5 | -0.0049 | <0.001 |
| 320 | ThalVPL | subcortical_ThalVPL_130 | +0.0044 | <0.001 |
| 325 | ThalMDm | subcortical_ThalMDm_135 | -0.0041 | <0.001 |
| 119 | DefaultA | LH_DefaultA_PFCm_1 | +0.0031 | <0.001 |
| 92 | ContA | LH_ContA_IPS_3 | -0.0027 | <0.001 |
| 112 | DefaultA | LH_DefaultA_PFCd_1 | -0.0023 | <0.001 |
| 192 | SomMotB | RH_SomMotB_S2_5 | -0.0022 | <0.001 |
| 322 | ThalIL | subcortical_ThalIL_132 | +0.0021 | <0.001 |
| 80 | LimbicB | LH_LimbicB_OFC_1 | +0.0019 | <0.001 |
| 235 | LimbicB | RH_LimbicB_OFC_2 | +0.0019 | <0.001 |
| 53 | DorsAttnA | LH_DorsAttnA_SPL_2 | +0.0018 | <0.001 |
| 159 | VisCent | RH_VisCent_ExStr_8 | -0.0016 | <0.001 |
| 304 | Amygdala | subcortical_Amygdala_46 | +0.0015 | <0.001 |
| 49 | DorsAttnA | LH_DorsAttnA_TempOcc_2 | +0.0012 | <0.001 |
| 262 | ContB | RH_ContB_PFClv_1 | -0.0011 | <0.001 |
| 141 | DefaultC | LH_DefaultC_Rsp_1 | +0.0007 | <0.001 |

#### 3.4. Classification following regression of age and sex effect

While in the main manuscript the effects of age and sex were included in classifier training along with network controllability metrics obtained for 2-back condition, here the classifier was trained using network controllability obtained for 2-back condition following regression of effects of age and sex. Supplementary Table 4 presents the classifier performance. Area under curve (AUC) and stability were marginally improved across all models when age and sex were regressed out prior to training rather than included as covariates. Combined model produced highest AUC and most stable (i.e. least variable accuracy) model.

| Supplementary Table 4 Classification performance summary for all models. All metrics are expressed in percentages (%). Values represent mean ± standard deviations across 5,000 evaluations (1,000 repetitions × 5-fold cross-validation). All models were trained following regression of age and sex from network controllability. | | | | |
| --- | --- | --- | --- | --- |
| **Metric** | **Average controllability** | **Modal controllability** | **Regional energy** | **Combined** |
| Accuracy | 79.4 ± 9.4 | 76.8 ± 9.2 | 75.9 ± 9.4 | 79.6 ± 8.5 |
| Balanced Accuracy | 78.8 ± 10.0 | 76.0 ± 9.6 | 74.7 ± 10.0 | 78.6 ± 9.1 |
| Sensitivity | 82.0 ± 12.1 | 80.4 ± 12.1 | 81.2 ± 12.0 | 84.0 ± 11.0 |
| Specificity | 75.5 ± 17.1 | 71.5 ± 16.9 | 68.2 ± 17.9 | 73.2 ± 16.0 |
| Precision | 84.1 ± 9.9 | 81.5 ± 9.5 | 79.9 ± 9.6 | 83.0 ± 8.9 |
| AUC | 87.3 ± 8.5 | 84.3 ± 8.7 | 85.0 ± 9.0 | 87.5 ± 7.6 |

#### 3.5. Additional predictive models - Using 2-back d-Prime and reaction time as response variables

Predictive modelling was repeated without principal component analysis, using only in-scanner 2-back d-Prime. All models achieved negative cross-validation R² (average controllability: R² = -0.091, p = 0.578; modal controllability: R² = -0.105, p = 0.822; activation energy: R² = -0.112, p = 0.882; combined: R² = -0.140, p = 0.876), indicating no generalisation above chance.

When 2-back response time was used as the response variable, all models achieved chance-level cross-validation performance (average controllability: R² = -0.052, p = 0.370; modal controllability: R² = -0.051, p = 0.365; activation energy: R² = -0.060, p = 0.549; combined: R² = -0.029, p = 0.222)

#### 3.6. Additional predictive models - effects of comorbidities and average motion

To investigate if comorbidities or average motion during scanning affected the models, predictive models were fitted again with these features as predictors along with controlability measures. Then, the distributions of R^2^ were compared with independent t-tests.

Adding binary comorbidity status (ADHD, Autism, dyslexia, sensory processing disorder) to the predictive models did not significantly affect predictive performance for any model (average controllability: t = -0.43, p = 0.669; modal controllability: t = -0.40, p = 0.692; activation energy: t = -0.35, p = 0.729; combined: t = -0.45, p = 0.651).

Similarly, adding mean framewise displacement as a covariate did not significantly affect predictive performance for any model (average controllability: t = -0.31, p = 0.755; modal controllability: t = -0.26, p = 0.792; activation energy: t = -0.25, p = 0.799; combined: t = -0.19, p = 0.850).

#### 3.7. Additional predictive models – unmedicated NF1 group

To examine whether predictive model performance was driven by medication effects, sensitivity analysis was conducted, where predictive models were estimated for the unmedicated NF1 group (n = 39).

In the final model, the PCA component explained 50.0% of variance in behavioural performance. Supplementary Table 5 presents the PCA coefficients of the first component, which were similar to the coefficients presented in the main manuscript. During cross-validation, the component explained 50.0% ± 4.1% of variance across 5,000 models (1,000 repetitions × 5-fold cross-validation).

| Supplementary Table 5 Coefficients of the first PCA component of working memory tasks in the unmedicated subsample (n = 39). Cross-validation values represent the mean ± standard deviation across 5,000 models (1,000 repetitions × 5-fold cross-validation). | | |
| --- | --- | --- |
| **Task** | **Final model coefficients** | **Cross-validation** |
| Verbal 0-back | 0.371 | 0.368 ± 0.031 |
| Verbal 2-back | 0.446 | 0.446 ± 0.013 |
| Visuospatial 0-back | 0.357 | 0.352 ± 0.044 |
| Visuospatial 1-back | 0.415 | 0.414 ± 0.022 |
| Visuospatial 2-back | 0.417 | 0.416 ± 0.018 |
| Visuospatial 3-back | 0.338 | 0.335 ± 0.029 |

Permutation testing revealed more accurate and more consistent predictive performance across controllability metrics in the unmedicated subsample only, compared to the full sample. Both the average controllability model (R² = 0.032 ± 0.022, null R² = −0.066 ± 0.075, p = 0.041) and the activation energy model (R² = 0.029 ± 0.022, null R² = −0.066 ± 0.075, p = 0.045) significantly exceeded the null distribution. The modal controllability model (R² = 0.025 ± 0.022, null R² = −0.066 ± 0.075, p = 0.054) and combined model (R² = 0.031 ± 0.023, null R² = −0.072 ± 0.082, p = 0.059) showed trends towards significance. Cross-validation Pearson's R values were consistently higher than in the full sample across all models (Supplementary Table 6), with the average controllability model improving from r = 0.223 to r = 0.302, and the activation energy model from r = 0.188 to r = 0.276.

| Supplementary Table 6 Behavioural prediction performance summary for the unmedicated subsample (n = 39). All models included age and sex as covariates. Values in cross-validation rows represent mean ± standard deviation across 1,000 repetitions. Final models were fitted to the full unmedicated subsample. The p-values of predictive performance were estimated based on permutation testing (10,000 permutations). | | | | |
| --- | --- | --- | --- | --- |
| **Metric** | **Average controllability** | **Modal controllability** | **Activation energy** | **Combined** |
| *Cross-validation* | | | | |
| *p-value* | **0.041** | 0.054 | **0.045** | 0.059 |
| R² | 0.032 ± 0.022 | 0.025 ± 0.022 | 0.029 ± 0.022 | 0.031 ± 0.023 |
| Pearson's R | 0.302 ± 0.137 | 0.246 ± 0.152 | 0.276 ± 0.149 | 0.242 ± 0.129 |
| ***Final model*** | | | | |
| R² | 0.039 | 0.032 | 0.038 | 0.095 |
| Pearson's R | 0.505 | 0.462 | 0.549 | 0.575 |

### 4. Additional covariates in group comparisons

To examine whether group differences in FC and network controllability were robust to medication-related and motion-related confounds, group comparisons of functional connectivity and network control theory, classifications and predictive modelling analyses were repeated with binary medication status and mean framewise displacement (FD) included as additional covariates alongside age and sex. Groups showed a trend towards an increased mean FD in NF1 sample (controls: 0.222 ± 0.205 cm, NF1: 0.300 ± 0.219 cm; t = -1.72, p = 0.090), warranting further investigation.

#### 4.1. Functional Connectivity

Most FDR-corrected FC group differences survived the inclusion of motion and medication as covariates. Specifically, the following key patterns were replicated: control network reductions involving left intraparietal sulcus and left lateral prefrontal cortex (control A); increased connectivity between left lateral ventral prefrontal cortex (control B) and bilateral medial prefrontal cortex (default A); default-limbic reductions between left ventral prefrontal cortex (default B) and bilateral orbital frontal cortex (limbic B); bilateral orbital frontal cortex reductions within the limbic network; and increased control-to-somatomotor connectivity between left lateral prefrontal cortex (control A) and right somatomotor cortex, and ventral attention-to-somatomotor connectivity between left insula (ventral attention B) and right somatomotor cortex.

The following connections emerged only in this analysis that controlled for motion and medication: increased connection between bilateral extrastriate cortex (central visual), increased connection between left dorsal prefrontal cortex (default B) and right medial posterior prefrontal cortex (ventral attention B), increased connection between left temporal cortex (control A) and left temporal cortex (default B), increased connection between left dorsal prefrontal cortex (default A) and right anterior temporal cortex (default B).

#### 4.2. Network Controllability

Group comparisons of network controllability repeated with medication and mean framewise displacement as additional covariates preserved the core pattern of disrupted controllability reported in the main manuscript, with a modestly expanded set of significant regions.

The dominant pattern of higher average controllability alongside lower modal controllability and lower activation energy in NF1 was replicated and extended across control, default, and dorsal attention networks. This pattern was preserved in left intraparietal sulcus (control A; ROI 92), bilateral lateral ventral prefrontal cortex (control B; ROIs 104, 262), bilateral precuneus (control C; ROIs 267, 268), bilateral superior parietal lobule (dorsal attention A; ROIs 53, 54), left frontal eye fields (dorsal attention B; ROI 63), bilateral medial prefrontal cortex (default A; ROI 119), left dorsal prefrontal cortex (default A; ROI 112), left lateral prefrontal cortex (default B; ROI 133), and bilateral retrosplenial cortex (default C; ROIs 141, 291).

Additional regions showing higher average controllability alongside lower modal controllability and lower activation energy in NF1 included right temporal cortex (default A; ROI 271), bilateral precuneus posterior cingulate cortex (default A; ROIs 118, 276), right ventral prefrontal cortex (default B; ROI 289), left precuneus (control C; ROI 109), left temporal pole (limbic A; ROI 85), bilateral orbital frontal cortex (limbic B; ROIs 80, 235, 237), bilateral somatomotor cortex (somatomotor A; ROIs 29, 34, 182). Right extrastriate cortex (central visual; ROI 159) continued to show higher average controllability alongside lower modal controllability, while a second extrastriate region (ROI 155) showed the reverse for average controllability only.

The opposite pattern of lower average controllability, higher modal controllability and higher activation energy was preserved in left temporal parietal cortex (ROIs 148, 149, 150) and right auditory cortex (somatomotor B; ROI 186). Additionally, right ventral posterior lateral thalamus (ROI 320) showed lower average controllability and lower modal controllability.

#### 4.3. Classification

Classification was repeated using network controllability measures obtained following regression of age, sex, medication status and mean framewise displacement. Supplementary Table 7 presents the classifier performance. All models showed reduced discriminative ability compared to both the main manuscript model and the demographics-only regression model, with AUC ranging from 72.2% to 80.8% compared to 82.7–85.5% in the main analysis. Average controllability again achieved the highest AUC (80.8%), while regional energy showed the greatest reduction in performance relative to the original analysis.

| Supplementary Table 7 Classification performance summary for all models. All metrics are expressed in percentages (%). Values represent mean ± standard deviations across 5,000 evaluations (1,000 repetitions × 5-fold cross-validation). All models were trained following regression of age and sex from network controllability. | | | | |
| --- | --- | --- | --- | --- |
| **Metric** | **Average controllability** | **Modal controllability** | **Regional energy** | **Combined** |
| Accuracy | 73.0 ± 10.3 | 71.9 ± 10.2 | 67.5 ± 10.2 | 71.3 ± 9.5 |
| Balanced Accuracy | 72.2 ± 10.6 | 70.6 ± 10.8 | 66.2 ± 10.7 | 70.4 ± 9.8 |
| Sensitivity | 76.4 ± 13.3 | 77.4 ± 13.1 | 73.0 ± 13.3 | 75.4 ± 13.1 |
| Specificity | 67.9 ± 16.9 | 63.8 ± 18.9 | 59.3 ± 18.9 | 65.4 ± 17.1 |
| Precision | 78.5 ± 10.0 | 76.8 ± 10.1 | 73.3 ± 9.9 | 77.0 ± 9.6 |
| AUC | 80.8 ± 10.1 | 79.5 ± 10.1 | 72.2 ± 10.9 | 79.7 ± 9.4 |

#### 4.4. Prediction

Predictive modelling was repeated with medication status and mean framewise displacement included as additional covariates, which showed improved performance relative to the main analysis (Supplementary Table 8). All three individual controllability models achieved significant cross-validation performance. Final model R² was improved across all models, with the combined model achieving the best fit (R² = 0.097). Framewise displacement was a significant negative predictor of working memory across all models.

| Supplementary Table 8 Behavioural prediction performance summary for models using network controllability measures with medication status and mean framewise displacement included as additional covariates. All metrics are expressed as proportions. Values in cross-validation rows represent mean ± standard deviations across 1,000 repetitions. Final models were fitted to the whole NF1 sample (n = 51). The p-values of predictive performance were estimated based on R² obtained during permutation testing | | | | |
| --- | --- | --- | --- | --- |
| **Metric** | **Average controllability** | **Modal controllability** | **Activation energy** | **Combined** |
| *Cross-validation* |  |  |  |  |
| *p*-value | **0.037** | **0.038** | **0.046** | 0.069 |
| R^2^ | 0.028 ± 0.017 | 0.028 ± 0.016 | 0.023 ± 0.017 | 0.022 ± 0.019 |
| Pearson's R | 0.281 ± 0.119 | 0.291 ± 0.121 | 0.240 ± 0.129 | 0.178 ± 0.113 |
| *Final model* |  |  |  |  |
| R^2^ | 0.046 | 0.044 | 0.038 | 0.097 |
| Pearson's R | 0.595 | 0.623 | 0.592 | 0.568 |

#### 4.5. Prediction beta weights

The covariate-extended predictive models largely preserved the two dominant patterns of brain-behaviour relationships identified in the main analysis (Supplementary Table 9). Additionally, left temporal pole (limbic A; ROI 85) emerged a new and strongest predictor across average and modal controllability models, where lower average controllability and higher modal controllability in left temporal pole were both significantly associated with better working memory performance (average controllability: β = -0.018, p < 0.001; modal controllability: β = +0.020, p < 0.001).

The first dominant pattern was preserved and higher working memory component scores in NF1 were predicted by lower average controllability, higher modal controllability, and higher activation energy in left intraparietal sulcus (control A; ROI 92), bilateral lateral ventral and temporal prefrontal cortex (control B; ROIs 104, 255, 258, 262), left superior parietal lobule and temporal occipital cortex (dorsal attention A; ROIs 49, 53, 54), bilateral orbital frontal cortex (limbic B; ROIs 80, 235), and left temporal parietal cortex (ROI 148). Left lateral prefrontal cortex (default B; ROI 133) showed the reverse of this pattern with higher values across all three measures predicting better performance, which was also consistent with the main manuscript.

Next, higher working memory component scores in NF1 were predicted by higher average controllability, lower modal controllability, and lower activation energy in left temporal parietal cortex (ROIs 149, 150), right auditory cortex (somatomotor B; ROI 186), right retrosplenial cortex (default C; ROI 291), right extrastriate cortex (central visual; ROI 159), right temporal cortex (default A; ROI 271), and right ventral posterior lateral thalamus (ROI 320), additionally left somatomotor cortex (ROI 34) continued this pattern but left somatomotor cortex (ROI 29) showed the opposite pattern across all three measures.

Consistently with the main manuscript, lower working memory component scores were predicted by with higher average controllability, higher modal controllability and higher activation energy in left dorsal and medial prefrontal cortex (default A; ROIs 112, 119).

| Supplementary Table 9 All beta weights from the final predictive models of behaviour (n = 51). Parcel indices, network labels, and full labels originate directly from the Schafer et al (2018) atlas. | | | | |
| --- | --- | --- | --- | --- |
| Parcel index | Network | Full label | β | p-value |
| ***Average controllability model*** | | | | |
| 85 | LimbicA | LH_LimbicA_TempPole_2 | -0.0176 | <0.001 |
| 104 | ContB | LH_ContB_PFClv_1 | -0.0119 | 0.004 |
| 54 | DorsAttnA | LH_DorsAttnA_SPL_3 | -0.0105 | 0.007 |
| 133 | DefaultB | LH_DefaultB_PFCl_1 | -0.0103 | 0.007 |
| 291 | DefaultC | RH_DefaultC_Rsp_1 | +0.0095 | 0.019 |
| 29 | SomMotA | LH_SomMotA_9 | +0.0092 | 0.018 |
| 253 | ContA | RH_ContA_PFCl_4 | -0.0092 | 0.017 |
| 149 | TempPar | LH_TempPar_4 | +0.0090 | 0.017 |
| 260 | ContB | RH_ContB_PFCld_2 | -0.0086 | 0.020 |
| 182 | SomMotA | RH_SomMotA_12 | +0.0077 | 0.041 |
| 276 | DefaultA | RH_DefaultA_pCunPCC_2 | +0.0073 | 0.057 |
| 34 | SomMotA | LH_SomMotA_14 | -0.0069 | 0.067 |
| 255 | ContB | RH_ContB_Temp_1 | -0.0061 | 0.083 |
| 289 | DefaultB | RH_DefaultB_PFCv_3 | +0.0059 | <0.001 |
| 150 | TempPar | LH_TempPar_5 | +0.0057 | <0.001 |
| 49 | DorsAttnA | LH_DorsAttnA_TempOcc_2 | -0.0057 | <0.001 |
| 186 | SomMotB | RH_SomMotB_Aud_1 | -0.0055 | <0.001 |
| 199 | DorsAttnA | RH_DorsAttnA_SPL_1 | -0.0055 | <0.001 |
| 53 | DorsAttnA | LH_DorsAttnA_SPL_2 | -0.0055 | <0.001 |
| 148 | TempPar | LH_TempPar_3 | +0.0053 | <0.001 |
| 320 | ThalVPL | subcortical_ThalVPL_130 | -0.0051 | <0.001 |
| 63 | DorsAttnB | LH_DorsAttnB_FEF_2 | -0.0046 | <0.001 |
| 159 | VisCent | RH_VisCent_ExStr_8 | -0.0042 | <0.001 |
| 271 | DefaultA | RH_DefaultA_Temp_1 | +0.0039 | <0.001 |
| 201 | DorsAttnA | RH_DorsAttnA_SPL_3 | -0.0035 | <0.001 |
| 237 | LimbicB | RH_LimbicB_OFC_4 | -0.0034 | <0.001 |
| 262 | ContB | RH_ContB_PFClv_1 | +0.0030 | <0.001 |
| 119 | DefaultA | LH_DefaultA_PFCm_1 | +0.0025 | <0.001 |
| 268 | ContC | RH_ContC_pCun_2 | +0.0024 | <0.001 |
| 235 | LimbicB | RH_LimbicB_OFC_2 | +0.0022 | <0.001 |
| 258 | ContB | RH_ContB_IPL_3 | -0.0022 | <0.001 |
| 267 | ContC | RH_ContC_pCun_1 | -0.0022 | <0.001 |
| 247 | ContA | RH_ContA_IPS_2 | -0.0021 | <0.001 |
| 92 | ContA | LH_ContA_IPS_3 | +0.0019 | <0.001 |
| 118 | DefaultA | LH_DefaultA_pCunPCC_5 | -0.0016 | <0.001 |
| 155 | VisCent | RH_VisCent_ExStr_5 | +0.0015 | <0.001 |
| 154 | VisCent | RH_VisCent_ExStr_4 | +0.0005 | <0.001 |
| 80 | LimbicB | LH_LimbicB_OFC_1 | +0.0005 | <0.001 |
| 112 | DefaultA | LH_DefaultA_PFCd_1 | -0.0004 | <0.001 |
| ***Modal controllability model*** | | | | |
| 85 | LimbicA | LH_LimbicA_TempPole_2 | +0.0198 | <0.001 |
| 104 | ContB | LH_ContB_PFClv_1 | +0.0117 | 0.003 |
| 54 | DorsAttnA | LH_DorsAttnA_SPL_3 | +0.0106 | 0.006 |
| 133 | DefaultB | LH_DefaultB_PFCl_1 | +0.0098 | 0.011 |
| 260 | ContB | RH_ContB_PFCld_2 | +0.0094 | 0.014 |
| 149 | TempPar | LH_TempPar_4 | -0.0089 | 0.020 |
| 291 | DefaultC | RH_DefaultC_Rsp_1 | -0.0085 | 0.029 |
| 29 | SomMotA | LH_SomMotA_9 | -0.0084 | 0.031 |
| 148 | TempPar | LH_TempPar_3 | -0.0077 | 0.042 |
| 276 | DefaultA | RH_DefaultA_pCunPCC_2 | -0.0073 | 0.047 |
| 182 | SomMotA | RH_SomMotA_12 | -0.0070 | 0.063 |
| 150 | TempPar | LH_TempPar_5 | -0.0067 | 0.070 |
| 255 | ContB | RH_ContB_Temp_1 | +0.0066 | 0.072 |
| 34 | SomMotA | LH_SomMotA_14 | +0.0061 | <0.001 |
| 271 | DefaultA | RH_DefaultA_Temp_1 | -0.0052 | <0.001 |
| 186 | SomMotB | RH_SomMotB_Aud_1 | +0.0050 | <0.001 |
| 199 | DorsAttnA | RH_DorsAttnA_SPL_1 | +0.0049 | <0.001 |
| 320 | ThalVPL | subcortical_ThalVPL_130 | +0.0048 | <0.001 |
| 125 | DefaultB | LH_DefaultB_Temp_4 | +0.0048 | <0.001 |
| 53 | DorsAttnA | LH_DorsAttnA_SPL_2 | +0.0047 | <0.001 |
| 49 | DorsAttnA | LH_DorsAttnA_TempOcc_2 | +0.0045 | <0.001 |
| 63 | DorsAttnB | LH_DorsAttnB_FEF_2 | +0.0038 | <0.001 |
| 237 | LimbicB | RH_LimbicB_OFC_4 | +0.0036 | <0.001 |
| 159 | VisCent | RH_VisCent_ExStr_8 | +0.0033 | <0.001 |
| 268 | ContC | RH_ContC_pCun_2 | -0.0033 | <0.001 |
| 92 | ContA | LH_ContA_IPS_3 | -0.0029 | <0.001 |
| 262 | ContB | RH_ContB_PFClv_1 | -0.0029 | <0.001 |
| 109 | ContC | LH_ContC_pCun_3 | -0.0026 | <0.001 |
| 201 | DorsAttnA | RH_DorsAttnA_SPL_3 | +0.0023 | <0.001 |
| 258 | ContB | RH_ContB_IPL_3 | +0.0016 | <0.001 |
| 155 | VisCent | RH_VisCent_ExStr_5 | -0.0015 | <0.001 |
| 112 | DefaultA | LH_DefaultA_PFCd_1 | -0.0014 | <0.001 |
| 267 | ContC | RH_ContC_pCun_1 | +0.0011 | <0.001 |
| 118 | DefaultA | LH_DefaultA_pCunPCC_5 | -0.0011 | <0.001 |
| 119 | DefaultA | LH_DefaultA_PFCm_1 | -0.0009 | <0.001 |
| 235 | LimbicB | RH_LimbicB_OFC_2 | -0.0004 | <0.001 |
| 80 | LimbicB | LH_LimbicB_OFC_1 | +0.0001 | <0.001 |
| ***Activation energy model*** | | | | |
| 104 | ContB | LH_ContB_PFClv_1 | +0.0110 | 0.005 |
| 148 | TempPar | LH_TempPar_3 | -0.0105 | 0.009 |
| 101 | ContB | LH_ContB_Temp_1 | +0.0103 | 0.010 |
| 54 | DorsAttnA | LH_DorsAttnA_SPL_3 | +0.0099 | 0.009 |
| 128 | DefaultB | LH_DefaultB_PFCd_1 | +0.0088 | 0.021 |
| 133 | DefaultB | LH_DefaultB_PFCl_1 | +0.0086 | 0.029 |
| 255 | ContB | RH_ContB_Temp_1 | +0.0080 | 0.032 |
| 48 | DorsAttnA | LH_DorsAttnA_TempOcc_1 | +0.0078 | 0.042 |
| 237 | LimbicB | RH_LimbicB_OFC_4 | +0.0075 | 0.046 |
| 245 | ContA | RH_ContA_Temp_1 | +0.0075 | 0.041 |
| 243 | LimbicA | RH_LimbicA_TempPole_5 | +0.0072 | 0.054 |
| 291 | DefaultC | RH_DefaultC_Rsp_1 | -0.0072 | 0.057 |
| 149 | TempPar | LH_TempPar_4 | -0.0071 | 0.057 |
| 29 | SomMotA | LH_SomMotA_9 | -0.0070 | <0.001 |
| 271 | DefaultA | RH_DefaultA_Temp_1 | -0.0065 | <0.001 |
| 182 | SomMotA | RH_SomMotA_12 | -0.0055 | <0.001 |
| 150 | TempPar | LH_TempPar_5 | -0.0053 | <0.001 |
| 109 | ContC | LH_ContC_pCun_3 | -0.0049 | <0.001 |
| 320 | ThalVPL | subcortical_ThalVPL_130 | +0.0047 | <0.001 |
| 92 | ContA | LH_ContA_IPS_3 | -0.0043 | <0.001 |
| 268 | ContC | RH_ContC_pCun_2 | -0.0041 | <0.001 |
| 186 | SomMotB | RH_SomMotB_Aud_1 | +0.0039 | <0.001 |
| 34 | SomMotA | LH_SomMotA_14 | +0.0035 | <0.001 |
| 119 | DefaultA | LH_DefaultA_PFCm_1 | +0.0034 | <0.001 |
| 112 | DefaultA | LH_DefaultA_PFCd_1 | -0.0030 | <0.001 |
| 63 | DorsAttnB | LH_DorsAttnB_FEF_2 | +0.0029 | <0.001 |
| 141 | DefaultC | LH_DefaultC_Rsp_1 | +0.0023 | <0.001 |
| 125 | DefaultB | LH_DefaultB_Temp_4 | +0.0020 | <0.001 |
| 235 | LimbicB | RH_LimbicB_OFC_2 | +0.0020 | <0.001 |
| 1 | VisCent | LH_VisCent_ExStr_1 | +0.0020 | <0.001 |
| 262 | ContB | RH_ContB_PFClv_1 | -0.0018 | <0.001 |
| 49 | DorsAttnA | LH_DorsAttnA_TempOcc_2 | +0.0017 | <0.001 |
| 80 | LimbicB | LH_LimbicB_OFC_1 | +0.0016 | <0.001 |
| 53 | DorsAttnA | LH_DorsAttnA_SPL_2 | +0.0013 | <0.001 |
| 159 | VisCent | RH_VisCent_ExStr_8 | +0.0010 | <0.001 |
| 55 | DorsAttnA | LH_DorsAttnA_SPL_4 | -0.0009 | <0.001 |
| 258 | ContB | RH_ContB_IPL_3 | +0.0007 | <0.001 |

#### 4.6. Using stable group differences in modelling

The analysis outlined in this Supplementary Material 4.2. demonstrated that some disruptions to network controllability measures in NF1 are robust and survive regardless of whether during group comparisons we control for motion or medication status or not. Therefore, these disruptions shared across the main manuscript and Supplementary Material 4.2., age, sex and medication were carried again to predictive modelling.

##### 4.6.1. Classification

Classification preserved good discriminative ability across all models (Supplementary Table 10). Average controllability again achieved the highest AUC (85.5%), matching the main manuscript performance, while modal controllability showed improvement relative to the covariate-extended analysis from Supplementary Table 4. (84.7% vs 79.5%).

| Supplementary Table 10. Classification performance summary for models using shared network controllability regions. All metrics are expressed in percentages (%). Values represent mean ± standard deviations across 5,000 evaluations (1,000 repetitions × 5-fold cross-validation). All models included age and sex as covariates. | | | | |
| --- | --- | --- | --- | --- |
| **Metric** | **Average controllability** | **Modal controllability** | **Regional energy** | **Combined** |
| Accuracy | 77.3 ± 9.9 | 76.2 ± 9.2 | 74.5 ± 9.6 | 72.3 ± 9.6 |
| Balanced Accuracy | 76.8 ± 10.5 | 75.7 ± 9.7 | 73.4 ± 10.3 | 71.2 ± 10.1 |
| Sensitivity | 79.1 ± 12.4 | 78.4 ± 12.6 | 78.9 ± 12.1 | 77.0 ± 12.6 |
| Specificity | 74.5 ± 18.3 | 73.0 ± 17.2 | 67.9 ± 18.3 | 65.5 ± 17.7 |
| Precision | 83.2 ± 10.7 | 82.1 ± 9.9 | 79.3 ± 10.0 | 77.5 ± 9.7 |
| AUC | 85.5 ± 8.9 | 84.7 ± 8.8 | 81.2 ± 10.1 | 79.4 ± 9.7 |

##### 4.6.2. Prediction

Predictive modelling using network controllability measures from regions showing stable group differences yielded modest generalisation performance (Supplementary Table 11). During cross-validation, all individual models showed low positive variance explained (average controllability: R² = 0.019 ± 0.018; modal controllability: R² = 0.017 ± 0.018; activation energy: R² = 0.019 ± 0.018), while the combined model performed least effectively (R² = 0.012 ± 0.018). None of the models significantly outperformed the null distribution.

In final models fitted to the full sample (n = 51), predictive performance improved slightly, with the combined model achieving the highest fit (R² = 0.061), followed by average controllability (R² = 0.029), activation energy (R² = 0.025), and modal controllability (R² = 0.024). Overall, restricting features to stable group-difference regions did not improve predictive performance relative to the main or covariate-extended analyses.

| Supplementary Table 11 Behavioural prediction performance summary for models using shared network controllability regions. All metrics are expressed as proportions. Values in cross-validation rows represent mean ± standard deviations across 1,000 repetitions. Final models were fitted to the whole NF1 sample (n = 51). The *p*-values of predictive performance were estimated based on R² obtained during permutation testing. | | | | |
| --- | --- | --- | --- | --- |
| **Metric** | **Average controllability** | **Modal controllability** | **Activation energy** | **Combined** |
| *Cross-validation* | | | | |
| *p*-value | 0.059 | 0.061 | 0.054 | 0.095 |
| R² | 0.019 ± 0.018 | 0.017 ± 0.018 | 0.019 ± 0.018 | 0.012 ± 0.018 |
| Pearson’s R | 0.199 ± 0.137 | 0.187 ± 0.140 | 0.198 ± 0.138 | 0.114 ± 0.123 |
| *Final model* | | | | |
| R² | 0.029 | 0.024 | 0.025 | 0.061 |
| Pearson’s R | 0.403 | 0.366 | 0.410 | 0.446 |
